# Nodosome inhibition as a novel broad-spectrum antiviral strategy against arboviruses and SARS-CoV-2

**DOI:** 10.1101/2020.11.05.370767

**Authors:** Daniel Limonta, Lovely Dyna-Dagman, William Branton, Tadashi Makio, Richard W. Wozniak, Christopher Power, Tom C. Hobman

**Author notes:** Address correspondence to Tom C. Hobman,.

## Abstract

In the present report, we describe two small molecules with broad-spectrum antiviral activity. These drugs block formation of the nodosome. The studies were prompted by the observation that infection of human fetal brain cells with Zika virus (ZIKV) induces expression of nucleotide-binding oligomerization domain-containing protein 2 (NOD2), a host factor that was found to promote ZIKV replication and spread. A drug that targets NOD2 was shown to have potent broad-spectrum antiviral activity against other flaviviruses, alphaviruses and SARS-CoV-2, the causative agent of COVID-19. Another drug that inhibits the receptor-interacting serine/threonine-protein kinase 2 (RIPK2) which functions downstream of NOD2, also decreased replication of these pathogenic RNA viruses. The broad-spectrum action of nodosome targeting drugs is mediated, at least in part, by enhancement of the interferon response. Together, these results suggest that further preclinical investigation of nodosome inhibitors as potential broad-spectrum antivirals is warranted.

## INTRODUCTION

Re-emerging and emerging RNA viruses represent a major threat to global public health. Vaccines and antiviral drugs which usually target a single virus species, are critical measures to prevent and control the spread of these pathogens. However, prophylactic or therapeutic drugs are not available for many of the most important RNA viruses in circulation today (1). While highly effective direct-acting antiviral drugs have been developed for a number of important human pathogens such as HIV-1, herpesvirus family members and hepatitis C virus (2), these drugs tend to be highly specific and are of limited use for treating other viral infections. In contrast, broad-spectrum antivirals would be expected to inhibit replication of multiple viruses, including emerging and re-emerging RNA viruses. Although several broad-spectrum antiviral compounds are in preclinical studies or in clinical trials, to date, no drug in this class has been licensed (3). Because hundreds of cellular factors are required for productive viral infection, targeting common host factors that are utilized by multiple viruses, may be a viable approach for developing broad-spectrum antivirals (4).

Herein we report the identification and characterization of a novel class of small molecules with broad-spectrum antiviral activity. These drugs selectively block the intracellular pattern recognition receptor NOD2 (nucleotide-binding oligomerization domain-containing protein 2), and a critical mediator of NOD2 signalling, RIPK2 (receptor-interacting serine/threonine-protein kinase 2). NOD2 recognizes the peptidoglycan muramyl dipeptide (MDP) that is found in bacterial cell walls, but it can also bind to viral RNA. In doing so, NOD2 induces formation of the nodosome and stimulates host defense against infections (5). Although NOD2 is important for the innate immune response against HIV-1 (6), cytomegalovirus (7) and syncytial respiratory virus (8), it has also been reported as a major pathogenic mediator of coxsackievirus B3-induced myocarditis (9).

Previously, we reported that Zika virus (ZIKV) infection of human fetal brain cells upregulates the expression of NOD2 (10) and here, we show that expression of this host protein promotes ZIKV replication. Using multiple human primary cell types, tissue explants, and cell lines, we found that the NOD2 blocking drug, GSK717, inhibits replication of flaviviruses, alphaviruses and SARS-CoV-2, the causative agent of COVID-19. The broad-spectrum activity of this drug is mediated in part by enhancement of the innate immune response. The RIPK2 blocking agent, GSK583, also potently inhibits these pathogenic RNA viruses. Together, data from our *in vitro* and *ex vivo* experiments suggest that nodosome inhibitors should be further investigated as broad-spectrum antivirals in preclinical studies.

## RESULTS

### ZIKV infection induces the inflammasome in primary human fetal brain cells

Previously, we reported that HFAs are likely the principal reservoirs for ZIKV infection and persistence in the human fetal brain (11). In a subsequent study (10), RNAseq analyses revealed that ZIKV infection of these cells upregulates multiple inflammasome genes including *GSDMD, IL-1 β, Casp1, NLRC5, GBP5*, and *NOD2*. In light of the recently identified links between the inflammasome and ZIKV neuropathogenesis (12, 13), we asked whether the activity of this multiprotein complex affected virus replication. Since our previous analysis (10) was performed using a strain of ZIKV not associated with microcephaly, we first confirmed that infection of HFAs with the pandemic ZIKV strain PRVABC-59 induced expression of multiple inflammasome genes. Indeed, *NOD2* and *GBP5* were upregulated more than 100-fold by ZIKV infection whereas other genes in this pathway were induced less than 50-fold (Fig. 1 A). Expression of inflammasome genes could also be induced by treatment of HFAs with human recombinant IFN-α, but less so than with the double-strand RNA mimic poly(I:C) (Fig. 1 B and C, Fig. S1 A-D). This may indicate that detection of viral RNA *per se* triggers inflammasome induction.

**Fig. 1.**
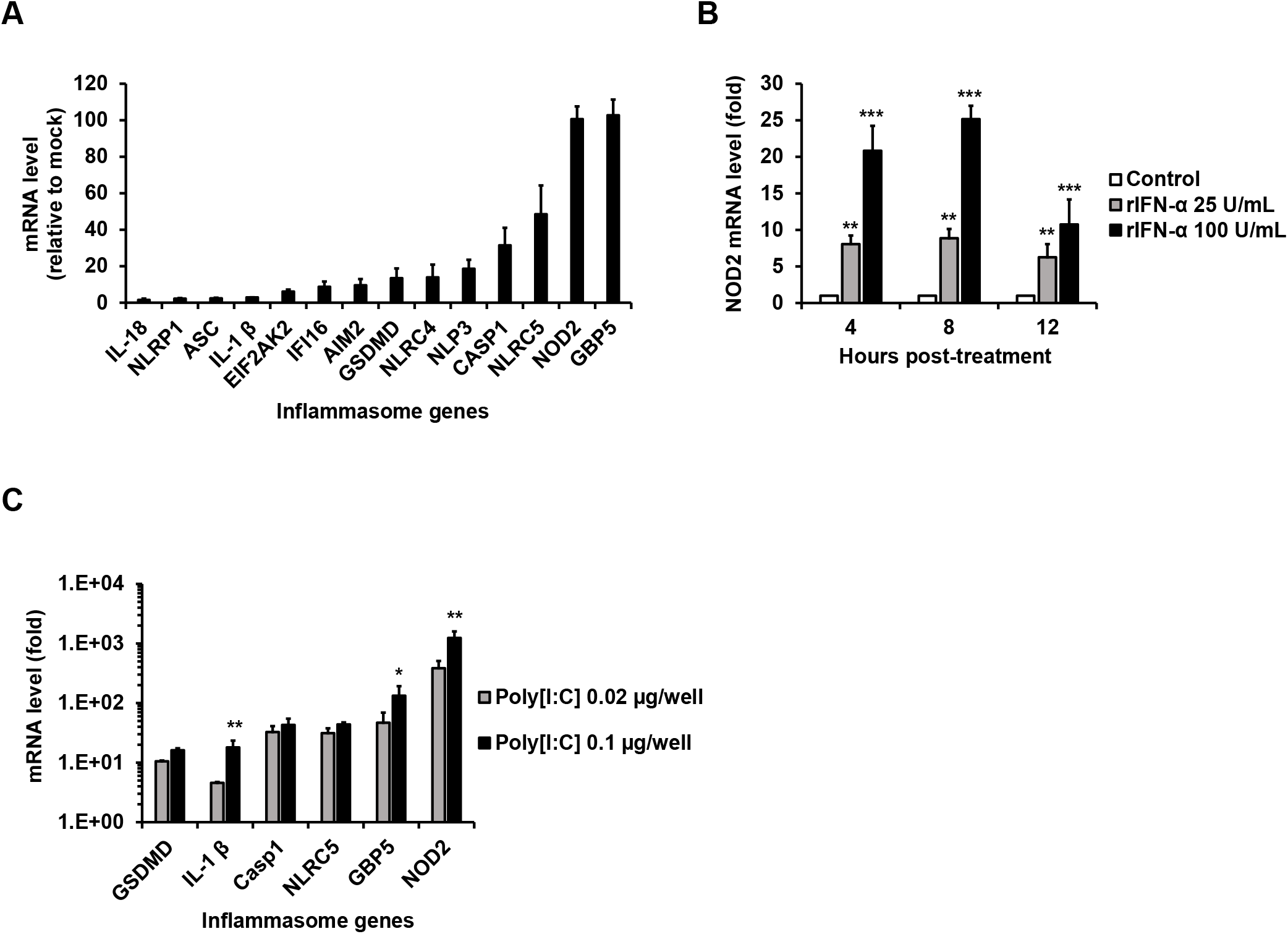
Inflammasome gene expression in HFAs is induced by ZIKV, IFN-α and Poly(I:C). (**A**) Relative inflammasome gene expression in HFAs infected with PRVABC-59 ZIKV (MOI=0.3) was determined by qRT-PCR 48-hours post-infection. (**B**) HFAs were treated with human recombinant IFN-α for 4, 8 and 12 hours after which relative *NOD2* expression was determined. (**C**) HFAs were transfected with Poly(I:C) for 12 hours after which relative inflammasome gene expression was determined. Error bars represent standard errors of the mean. \**P* < 0.05, \*\**P* < 0.01, and \*\*\**P* < 0.001, by the Student *t* test.

### NOD2 expression promotes ZIKV multiplication by suppression of the innate immune response in HFAs

As *NOD2* was one of the most upregulated inflammasome genes, we examined how reduced NOD2 expression in HFAs affected replication of the PRVABC-59 strain of ZIKV. Compared to cells transfected with a non-targeting siRNA, replication of ZIKV in HFAs transfected with NOD2-specific siRNAs was significantly reduced (Fig. 2 A and B).

**Fig. 2.**
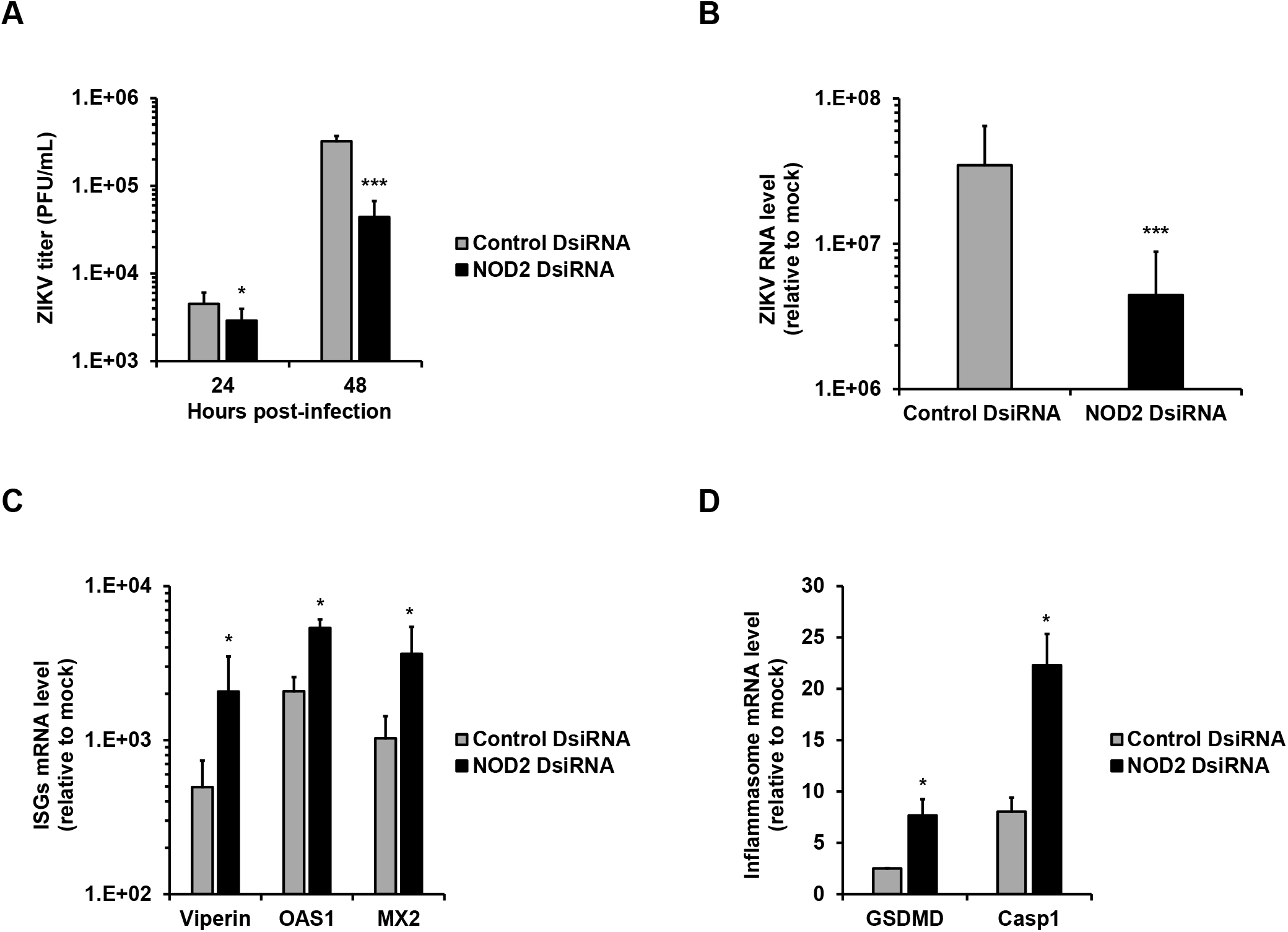
NOD2 silencing suppresses ZIKV multiplication and enhances the expression of interferon-stimulated and inflammasome genes. HFAs were transfected with NOD2-specific or non-silencing siRNAs for 24-hours and then infected with ZIKV (MOI=0.05). Cell culture media or total cellular RNA were harvested after 24- and 48-hours for plaque assay (**A**) or qRT-PCR at 48-hours post-infection. Relative levels of viral genome (**B**), and interferon-stimulated genes (**C**) *Viperin*, 2’-5’-oligoadenylate synthetase 1 (*OAS1*) and Myxovirus resistance protein 2 (*MX2*) as well as inflammasome genes (**D**) gasdermin D (*GSDMD*) and Caspase 1 (*Casp1*) are shown. Values are expressed as the mean of three independent experiments. Error bars represent standard errors of the mean. \**P* < 0.05, and \*\*\**P* < 0.001, by the Student *t* test.

Recently, we reported that ZIKV-induced expression of fibroblast growth factor 2 in fetal brain increases viral replication by inhibiting the interferon response (10). As NOD2 is an intracellular pattern recognition receptor that recognizes bacterial MDP as well as viral RNA (5), we questioned whether the effect of NOD2 expression on viral replication was related to its actions on the innate immune response. To address this, relative expression of ISGs was determined in ZIKV-infected HFAs after *NOD2* silencing. *NOD2* knockdown was associated with significant upregulation of several important ISGs including *Viperin, OAS1*, and *MX2* (Fig. 2 C) as well as the prototypic inflammasome genes *GSDMD* and *Casp1* (Fig. 2 D). Of note, *NOD2* silencing reduced *NOD2* mRNA expression and was not cytotoxic for HFAs (Fig. S1 E and F).

### ZIKV replication and spread are inhibited by blocking NOD2 function in fetal brain

A number of specific NOD2 inhibitors have been developed to treat inflammatory diseases (14). To determine if these drugs have antiviral activity, HFAs infected with ZIKV were treated with or without subcytotoxic concentrations of the anti-NOD2 compound, GSK717, for up to 72 hours. GSK717 reduced viral titers in a concentration-dependent manner regardless of whether cells were infected at low or high MOI (Fig. 3 A and B).

**Fig. 3.**
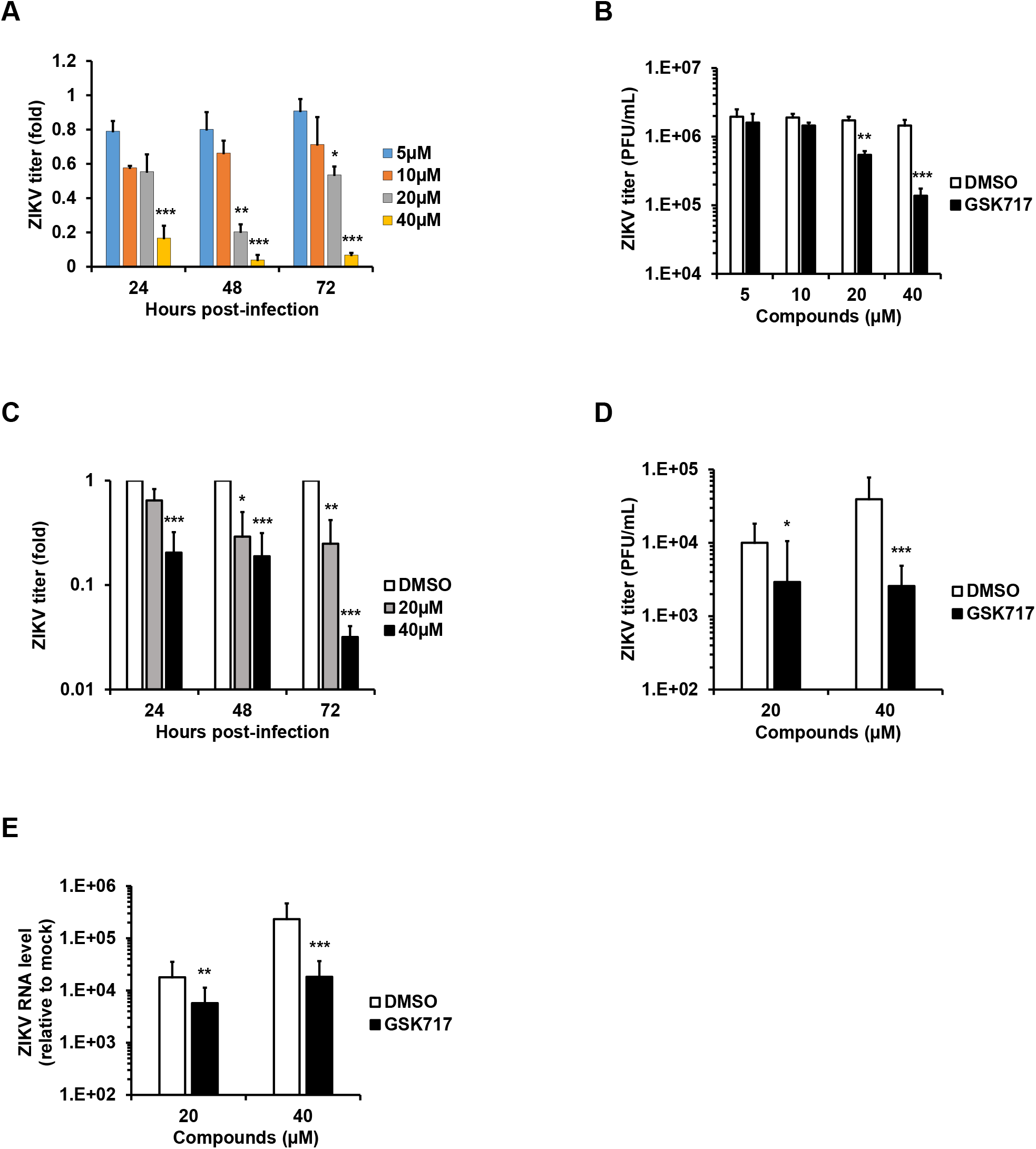
The anti-NOD2 drug GSK717 inhibits ZIKV replication. ZIKV-infected HFAs (MOI=0.05-5) were treated with DMSO or NOD2 blocking agent GSK717 after which viral titers were determined daily for up to 72-hours post-infection. ZIKV titers as relative fold (MOI=0.05) for 72 hours (**A**) and as PFU/mL at 48 hours post-infection (MOI=5) (**B**) are shown. Explants of human fetal brain (15-19 week old donors) were infected with the microcephalic ZIKV strain PRVABC-59 followed by GSK717 or DMSO treatment. Viral titers are shown as relative fold for 72 hours (**C**) and as PFU/mL at 48 hours post-infection (**D**). At 72 hours, the explant tissue was collected for viral RNA quantitation by qRT-PCR (**E**). Values are expressed as the mean of three independent experiments. Error bars represent standard errors of the mean. \**P* < 0.05, \*\**P* < 0.01, and \*\*\**P* < 0.001, by the Student *t* test.

We also investigated whether GSK717 could block ZIKV replication in explanted human fetal brain tissue as described previously (15). Data in Fig. 3 C-E show that GSK717 treatment significantly inhibited replication of viral genomic RNA and reduced viral titers of ZIKV by as much as 33-fold. Similar results were observed in human primary embryonic pulmonary fibroblasts (HEL-18) thus demonstrating that the antiviral effect of GSK717 is not limited to fetal brain tissue (Fig. S2 A-C).

### NOD2 inhibitor blocks ZIKV infection and spread in multiple cell lines

We next examined whether GSK717 also inhibits ZIKV replication in human non-prenatal cell types, including cell lines usually used for anti-flavivirus drug screening such as A549 (pulmonary) and Huh7 (hepatoma) cells (16) as well as the astrocytoma cell line U251. GSK717 significantly reduced ZIKV titers in these human cell lines in a dose-dependent manner regardless of whether low (0.05) and high (5) MOI were used for infection (Fig. S2 D-F). GSK717 also showed a concentration-dependent inhibitory effect on ZIKV replication in mouse embryonic fibroblasts (data not shown). Consistent with its ability to reduce viral titers, treatment with GSK717 significantly reduced the numbers of infected cells in A549 cultures (Fig. 4 A and B). Interestingly, GSK717 inhibited ZIKV replication even when added 12-or 24-hours post-infection (Fig. S2 G). None of the GSK717 concentrations used in the cell-based assays were cytotoxic at the examined time points (Fig. S3).

**Fig. 4.**
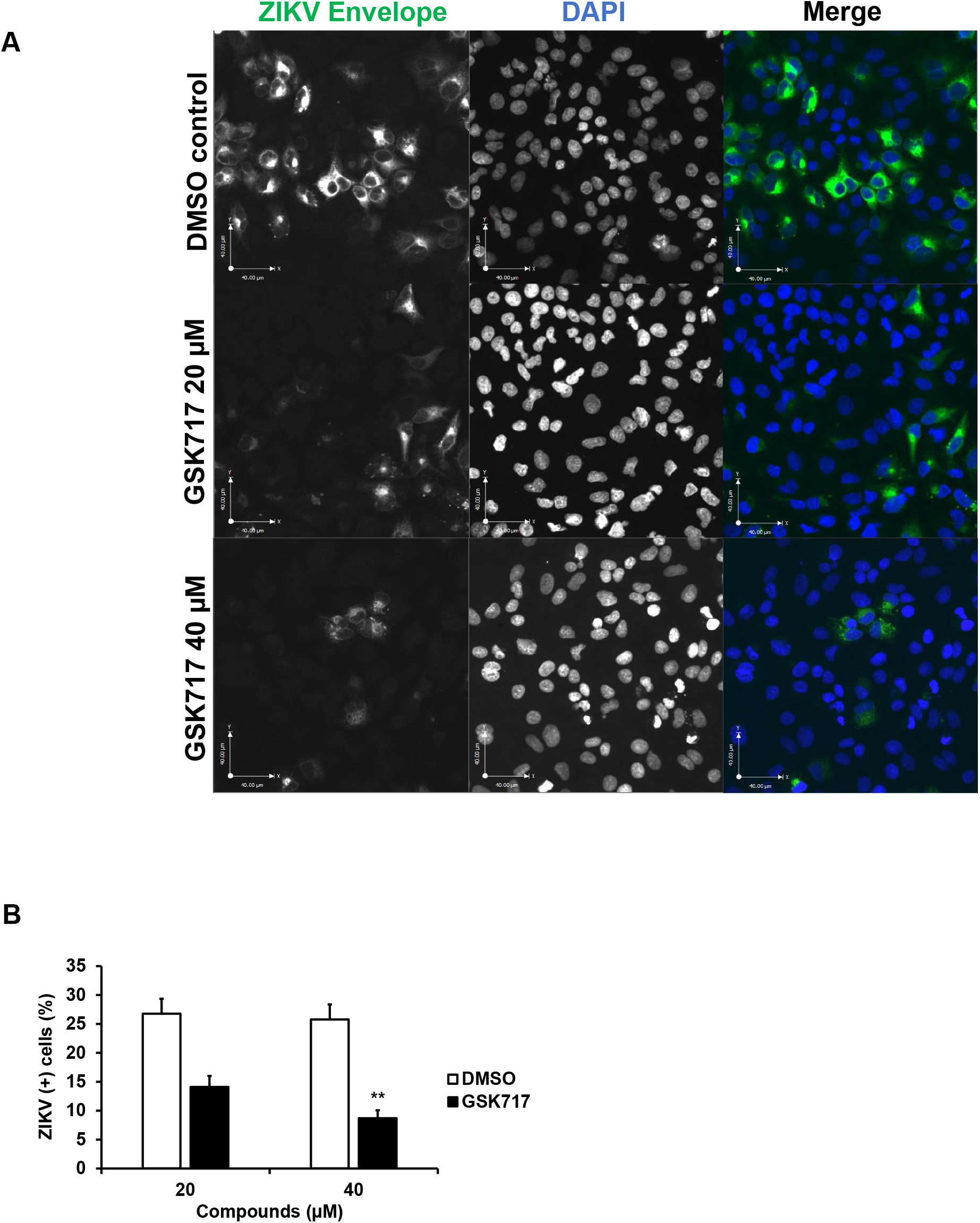
The anti-NOD2 drug GSK717 blocks the spread of ZIKV infection. (**A**) Representative confocal imaging (20X) showing antiviral effect of GSK717 at 20 μM and 40 μM. A549 cells were infected with ZIKV (MOI=1) followed by treatment with DMSO or GSK717 at 20 or 40 μM for 48 hours before processing for indirect immunofluorescence. ZIKV-infected cells were identified using a mouse monoclonal antibody (4G2) to envelope protein and Alexa Fluor 488 donkey anti-mouse to detect the primary antibody. Nuclei were stained with DAPI. Images were acquired using a spinning disk confocal microscope equipped with Volocity 6.2.1 software. (**B**) Infected cells in 10 different fields from samples treated with GSK717 or DMSO were counted. Values are expressed as the mean of three independent experiments. Error bars represent standard errors of the mean. \*\**P* < 0.01, by the Student *t* test.

### DENV replication is inhibited by the anti-NOD2 drug GSK717

To determine if GSK717 could inhibit replication of other flaviviruses, we next focused on DENV, the most important arbovirus in terms of morbidity and mortality and the causative agent of Dengue Hemorrhagic Fever/Dengue Shock Syndrome (17). A549 cells infected with DENV-2 strain 16681 (18) were treated with or without increasing concentrations of GSK717. Data in Fig. 5 A-C show that GSK717 reduced DENV titers by >90% when used at 20-40 μM. Blocking NOD2 function with GSK717 also dramatically reduced the number of viral antigen positive-A549 cells after 48 hours of infection (Fig. S4 A and B).

**Fig. 5.**
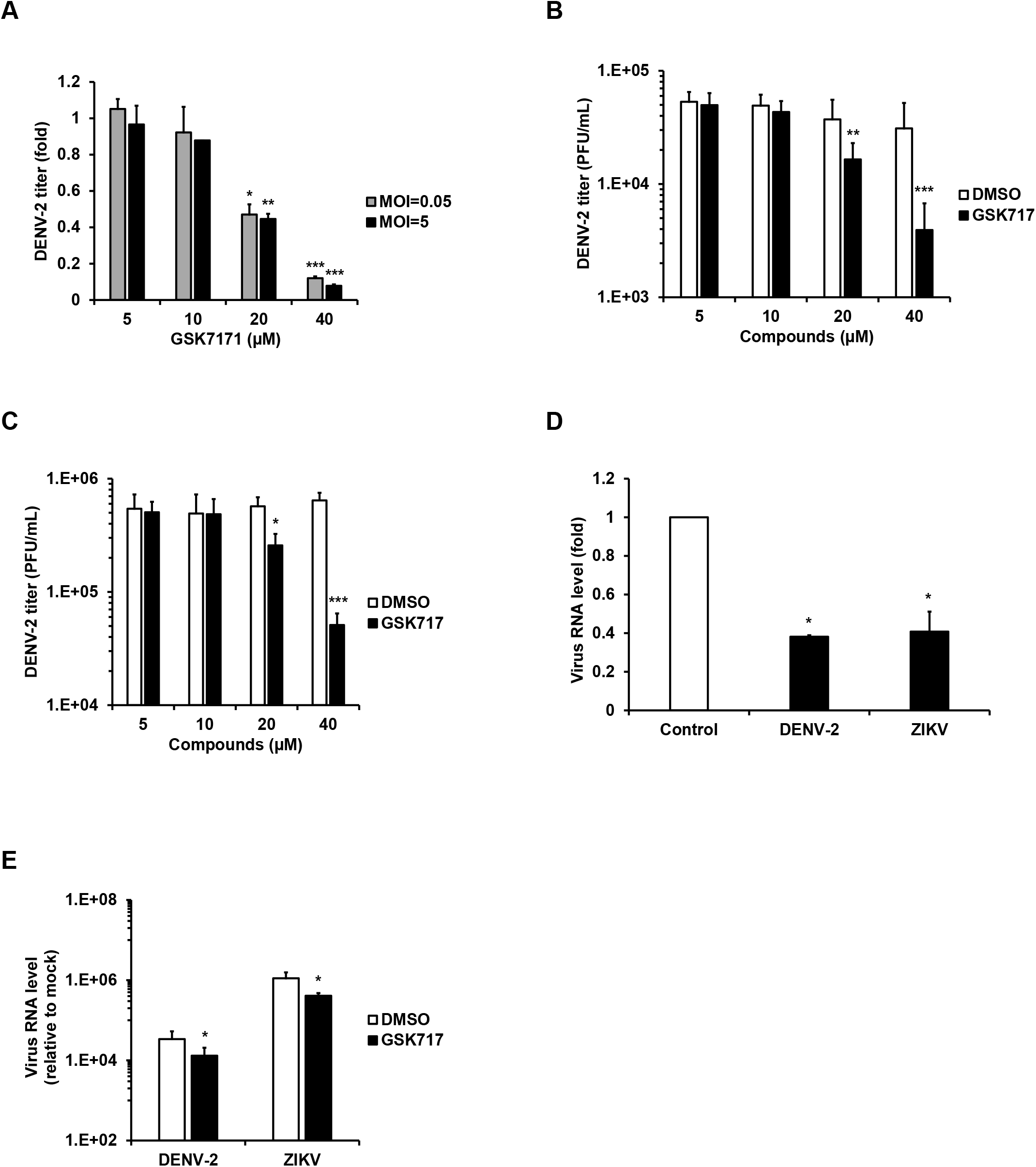
The anti-NOD2 drug GSK717 inhibits DENV replication. A549 cells were infected with DENV-2 (MOI=0.05-5) and treated with GSK717 or DMSO for 48-hours after which cell culture media were harvested for plaque assay. Viral titers as relative fold using MOI of 0.05 or 5 (**A**) and as PFU/mL using MOI of 0.05 (**B**) or 5 (**C**) are shown. Cells were co-infected with DENV-2 and ZIKV (MOI=0.1) followed by GSK717 or DMSO treatment for 48 hours before collecting total cellular RNA for qRT-PCR. Viral RNA levels as relative fold (**D**) and as relative to mock (**E**) are presented. Values are expressed as the mean of three independent experiments. Error bars represent standard errors of the mean. \**P* < 0.05, \*\**P* < 0.01, and \*\*\**P* < 0.001, by the Student *t* test.

Finally, as co-infections of DENV-2 with ZIKV have been reported in Latin American (19, 20) and South Asian countries (21), we assessed how GSK717 affected virus replication in A549 simultaneously infected with ZIKV and DENV-2. Results from qRT-PCR analyses revealed that levels of both ZIKV and DENV genomic RNA were reduced by ~60% in GSK717-treated cells (Fig. 5 D and E).

### NOD2 function is important for replication of alphaviruses and coronaviruses

Nodosome formation can be induced following infection by multiple types of RNA viruses (22). As such, we next determined whether GSK717 could inhibit replication of the mosquito-transmitted alphavirus, MAYV. The recent outbreak strain TRVL 15537 MAYV was used for these experiments. Supernatants from infected A549 cell cultures treated with or without GSK717 were collected for viral titer determination at 48-hours post-infection after which plaque assays were performed. Similar to what was observed in flavivirus-infected cells, we found that GSK717 reduced MAYV titers by as much as 95% (Fig. 6 A and B). MAYV and DENV-2 co-circulate in the same areas of South America and dual infections have been reported recently (23). Treatment of co-infected A549 cells with GSK717 resulted in significant inhibition of both DENV-2 and MAYV although replication of the latter was affected to a larger degree (Fig. 6 C and D).

**Fig. 6.**
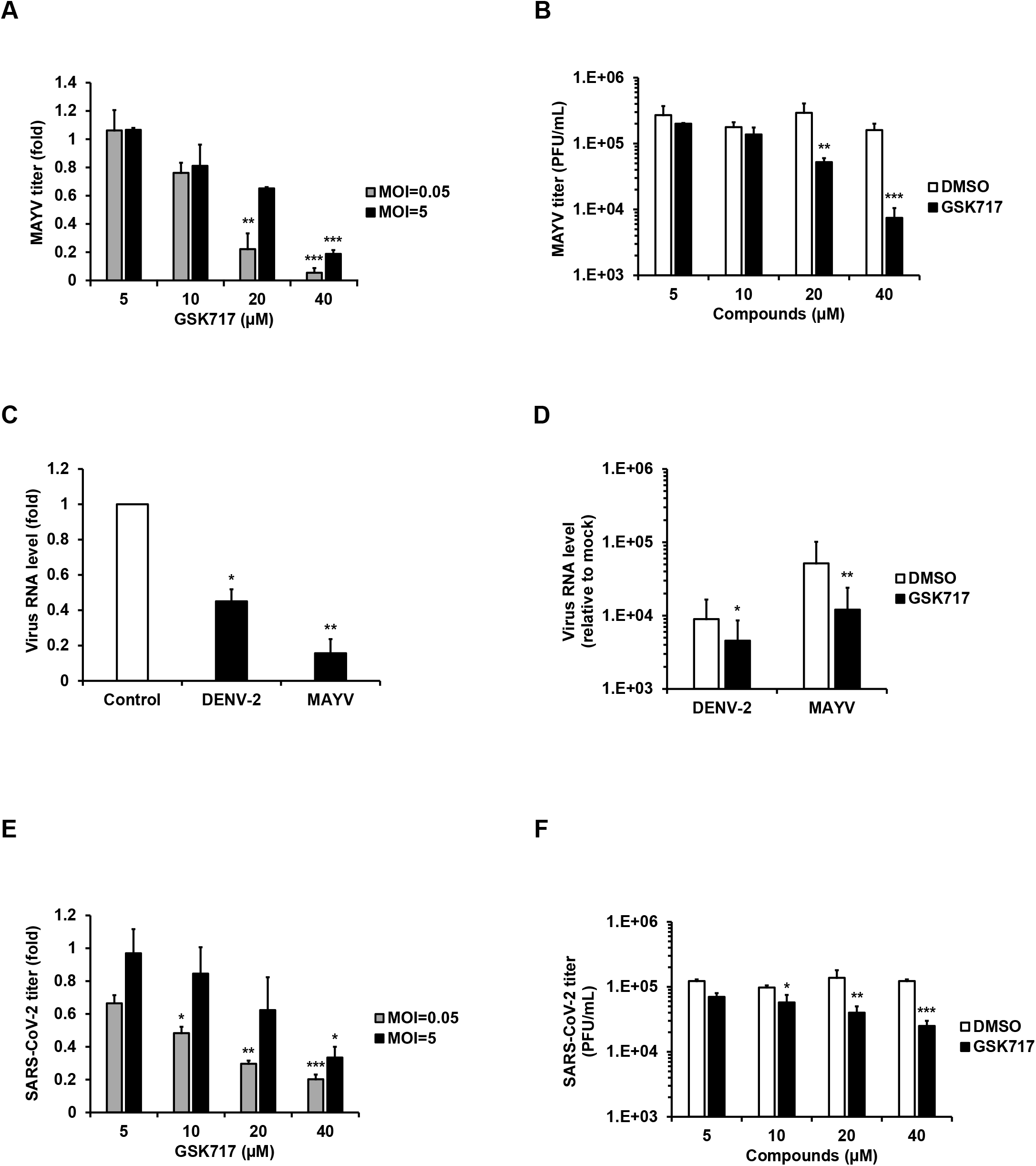
The anti-NOD2 drug GSK717 inhibits replication of MAYV and SARS-CoV-2 infection. A549 cells infected with low (0.05) and high (5) MOI of MAYV were treated with GSK717 or DMSO for 48 hours followed by collection of supernatants for plaque assay. Relative (**A**) and absolute (**B**) viral titers are shown with both MOI and with MOI of 0.05 respectively. Cells co-infected with MAYV and DENV-2 (MOI=0.1) were treated for 48 hours with GSK717 or DMSO before total cellular RNA was collected for qRT-PCR. Viral RNA data as relative fold (**C**) and as relative to mock (**D**) are shown. ACE2-SK-N-SH were infected with SARS-CoV-2 (MOI=0.05-5) and treated with GSK717 or DMSO for 48 hours after which culture supernatants were harvested for plaque assay. Viral titers as relative fold (**E**) and as PFU/mL (**F**) are shown using both MOI and the MOI of 0.05 respectively. Values are expressed as the mean of three independent experiments. Error bars represent standard errors of the mean. \**P* < 0.05, \*\**P* < 0.01, and \*\*\**P* < 0.001, by the Student *t* test.

Because SARS-CoV-2 infection reportedly activates the inflammasome *in vitro* (24) and in patients (25, 26), we tested how NOD2 inhibition affected replication of this pandemic coronavirus. A neuroblastoma cell line (SK-N-SH) stably expressing ACE2 was infected with a Canadian isolate of SARS-CoV-2 and treated with or without GSK717. At 48-hours post-infection, culture supernatants were collected for viral titer determination by plaque assay using Vero-E6 cells. Data in Fig. 6 E and F show that pharmacological inhibition of NOD2 resulted in reduction of SARS-CoV-2 titers similar to what was observed with arboviruses.

### Inhibition of downstream RIPK2 supresses replication of multiple RNA viruses including SARS-CoV-2

RIPK2 is a critical mediator of NOD2 signalling. Binding of MDP to NOD2 leads to self-oligomerization of NOD2 molecules, followed by homotypic interactions between the C-terminal caspase activation and recruitment domain of NOD2 and RIPK2. This results in the activation of transcription factors that drive the expression of multiple proinflammatory cytokines, chemokines and anti-bacterial proteins (27).

Although RIPK2 mRNA was not upregulated in ZIKV infected-HFAs (10), we questioned whether pharmacological inhibition of this protein would also reduce replication of arboviruses such ZIKV, DENV-2 and MAYV. Infected A549 cells were treated with or without GSK583, a highly potent and selective inhibitor of the NOD2 binding domain of RIPK2 (28). Significant reduction in viral titers was observed in GSK583-treated cells infected with ZIKV, DENV-2 or MAYV at 12- and 24-hours post-infection (Fig. 7 A-C, Fig. S5 A-D). The antiviral action of GSK583 was corroborated in another human cell line, Huh7, infected with MAYV (MOI=0.1) (data not shown).

**Fig. 7.**
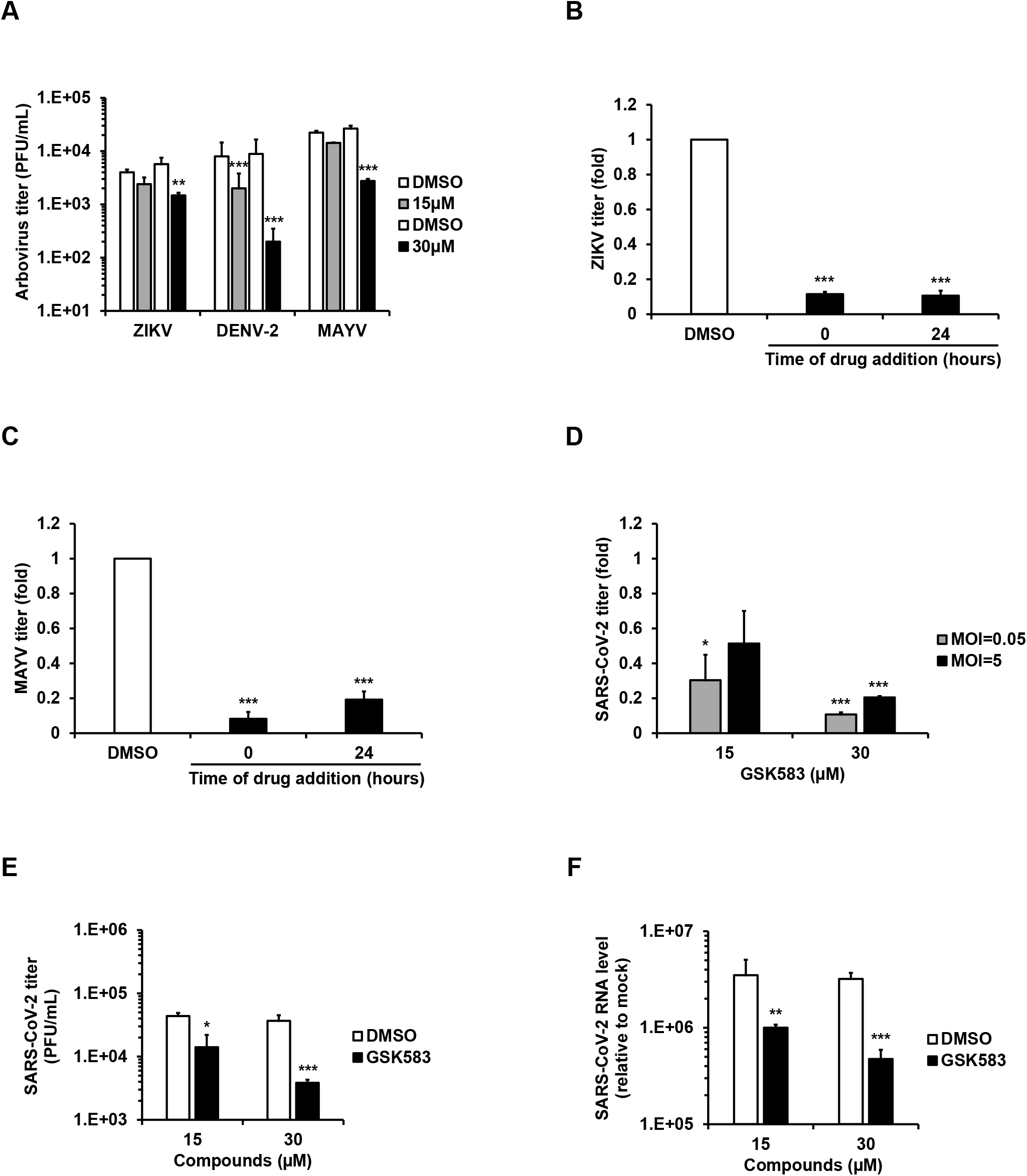
The anti-RIPK2 drug GSK583 has broad-spectrum antiviral activity. (**A**) A549 cells infected separately with three different arboviruses (MOI=0.1) were treated with the RIPK2 inhibitor GSK583 or DMSO alone for 24 hours followed by supernatant collection for plaque assay. Arbovirus titers as relative fold is presented. Cells were infected with ZIKV (**B**) or MAYV (**C)** at the MOI of 0.1 followed by the addition of GSK583 or DMSO immediately after infection (0 hours) or 24 hours post-infection. Viral titers, shown as relative fold, in supernatants were determined 24 hours after the treatment. ACE2-SK-N-SH cells infected with SARS-CoV-2 at low (0.05) and high (5) MOI were treated with GSK583 or DMSO for 24 hours followed by supernatant and total cellular RNA collection. Viral titers using both MOI as relative fold (**D**) and using MOI of 0.05 as PFU/mL (**E**) are shown. Relative viral RNA to mock level (**F**) with the MOI of 0.05 is shown. Values are expressed as the mean of three independent experiments. Error bars represent standard errors of the mean. \**P* < 0.05, \*\**P* < 0.01, and \*\*\**P* < 0.001, by the Student *t* test.

Similarly, indirect immunofluorescence microscopy analyses confirmed that inhibition of RIPK2 reduced the number of viral antigen-positive cells in ZIKV, DENV-2 and MAYV-infected A549 cultures (Fig. S6 A-D).

We tested the effect of GSK583 on replication of SARS-CoV-2 in ACE2-SK-N-SK cells. At 12 and 24-hours post-infection, culture supernatants and cell lysates were collected for viral titer determination by plaque assay and viral ARN quantification using qRT-PCR. A significant concentration-dependent reduction in viral multiplication was observed in GSK583-treated cells (Fig. 7 D-F and Fig. S7 A). A time-of-addition assay (drug treatment at 0- and 24-hour post-infection) demonstrated that the RIPK2 inhibitor was able to reduce virus replication even when the drug was added well after viral infection had occurred (Fig. S7 B). Quantitation of infection by indirect immunofluorescence showed that GSK583 treatment reduced the numbers of infected cells in a monolayer culture (Fig. S7 C and D).

Finally, when RIPK2 inhibition experiments were conducted using Calu-3 and Huh7 cells for 24 hours, ~60% and ~90% reduction in titers respectively were observed with the highest concentration of RIPK2 inhibitor (Fig. S8 A and B). No cytotoxic effect of GSK583 were detected in the cell lines used for coronavirus assays (Fig. S8 C-E).

## DISCUSSION

ZIKV co-circulates in some of the same endemic regions as other arboviruses including chikungunya virus, DENV, and MAYV. Symptoms of acute infection caused by these arboviruses such as fever, rash, joint pain, and ocular manifestations are common which complicates clinical diagnosis of mono- and co-infections (17, 29, 30). Differential serological diagnosis is further hindered by flavivirus antigen cross-reactivity (17, 30). Given the issues with clinical/laboratory diagnosis and lack of effective vaccines against most arboviruses, development of broad-spectrum antivirals against these pathogens should be a high priority.

The ongoing pandemic caused by SARS-CoV-2 poses a different set of challenges. Despite concerted efforts to repurpose and find new antiviral drugs (31–33), so far only remdesivir has shown modest efficacy in the acute stages of COVID-19 (34, 35). While more than 200 SARS-CoV-2 vaccine candidates are in accelerated development at preclinical and clinical stages (36), it will likely take another year or more before they are broadly available to the general population as safety and efficacy still need to be evaluated (37, 38).

In this study, we characterized the broad-spectrum antiviral activities of nodosome inhibitors GSK717 and GSK583. These small molecules display robust antiviral action against multiple RNA viruses and may hold promise as pan-flavivirus inhibitors. First, we showed that NOD2 expression promotes ZIKV multiplication in HFAs which are the main target of this flavivirus in the fetal brain (10, 11). Next, we demonstrated that the NOD2 inhibitor GSK717 blocks infection by and spread of ZIKV in human fetal brain and cell lines. NOD2 inhibition also reduced replication of the related DENV, the alphavirus MAYV and the pandemic coronavirus SARS-CoV-2. Blocking the NOD2 downstream signaling kinase RIPK2 with GSK583 (which does not affect its catalytic activity) significantly inhibited replication of these viral pathogens.

Gefitinib is an FDA-approved drug for treatment of lung, breast and other cancers. It works by reducing the activity of the epidermal growth factor receptor (EGFR) tyrosine kinase domains. Of note, this drug also inhibits the tyrosine kinase activity of RIPK2 (39) and has been shown to inhibit replication of DENV and release of pro-inflammatory cytokines from infected human primary monocytes (40). The authors suggested a role for EGFR/RIPK2 in DENV pathogenesis and that gefitinib may be beneficial in the treatment of dengue patients. Similarly, the work here which demonstrated the antiviral activity of NOD2 and RIPK2 inhibitors using tissue explants, primary cells and cell lines, support the potential clinical use of these compounds in mono or co-infections by arboviruses as well as coronavirus infections at early and/or advanced stages.

As GSK717 and GSK583 were developed primarily for immune-mediated inflammatory conditions, their anti-inflammatory effects may have the added benefit of reducing the hyperinflammatory state associated with flavi-, alpha- and coronavirus diseases (17, 29, 30, 41). Finally, our findings raise potential concerns regarding adjuvants in viral vaccines that augment NOD2 as an immune strategy (42, 43) since this immune signaling protein is not a restriction factor but rather an enhancement factor for multiple pathogenic RNA viruses.

The current study illustrates how the identification of a drug target through transcriptomic analyses of virus-infected cells can lead to novel broad-acting host-directed antiviral strategies with a high barrier of resistance. Increased NOD2 expression may be a novel mechanism of immune evasion that viruses use to evade the innate immune response. Conversely, drugs that block nodosome formation appear to have broad-spectrum antiviral activity by enhancing the interferon response. Collectively, our results warrant consideration of these and related compounds as broad-spectrum antiviral drug candidates for further preclinical development.

## MATERIALS AND METHODS

### Ethical Approval

Human fetal brain tissues were obtained from 15-19-week aborted fetuses with written consent from the donor parents and prior approval under protocol 1420 (University of Alberta Human Research Ethics Board).

### Cells, explant cultures and viruses

ZIKV (PRVABC-59), Dengue virus (DENV-2, 16681), and Mayaro virus (MAYV, TRVL 15537) were propagated in *Aedes albopictus* C6/36 cells grown in Minimum Essential Medium (MEM, Thermo Fisher Scientific, Waltham, MA). SARS-CoV-2 (SARS-CoV-2/CANADA/VIDO 01/2020) was propagated in Vero-E6 cells grown in Dulbecco’s Modified Eagle Medium (DMEM, Thermo Fisher Scientific). A549, Huh7, U251, Vero (ATCC, Manassas, VA) and ACE2-hyperexpressing SK-N-SH cells were maintained in DMEM while HEL-18 human primary embryonic pulmonary fibroblasts and Calu-3 cells (ATCC) were maintained in Roswell Park Memorial Institute 1640 medium (RPMI, Thermo Fisher Scientific) and MEM respectively. Human fetal astrocytes (HFAs) and fetal brain tissue explants were prepared from multiple donations (n=8), as described previously (15). For infection, cells or tissue explants were incubated with virus (MOI 0.05-5) for 1-2 hr or overnight respectively at 37°C using fresh media supplemented with fetal bovine serum (Thermo Fisher Scientific). For co-infection assays, A549 cells were infected simultaneously with DENV-2 and ZIKV or MAYV at an MOI of 0.1 for 3 hours. Culture of cells, tissue explants, construction of the ACE2-SK-N-SH cells, and viral infections are described in more detail in Supplemental Material.

### qRT-PCR

RNA from cells and tissue was extracted using NucleoSpin RNA (Macherey-Nagel GmbH & Co, Düren, Germany) kits. Real-time qRT-PCR was performed in a CFX96 Touch Real-Time PCR Detection System instrument (Bio-Rad, Hercules, CA) using ImProm-II Reverse Transcriptase (Promega, Madison, WI). For more details about the protocols, and primers used in this work please see Supplemental Material.

### Poly(I:C) transfection

HFAs grown in 96-well plates (Greiner, Kremsmünster, Austria) were transfected with polyinosinic:polycytidylic acid (Poly(I:C) (Sigma-Aldrich, St. Louis, MO) at a concentration of 0.02 or 0.1 μg/well using TransIT (0.3 μL/well, Mirus Bio LLC, Madison, WI). At 12 hours post-transfection, total RNA was extracted and transcripts levels for IFN-stimulated genes (ISGs) were quantified by qRT-PCR.

### Human recombinant INF-α assay

HFAs in 96-well plates (Greiner) were treated with or without 25-100 U/mL of human recombinant IFN-α (Sigma-Aldrich) for 4-12 hours after which total RNA was isolated and subjected to qRT-PCR in order to measure expression of ISGs.

### Viral titer assay

Titers were determined in Vero CCL-81 and Vero-E6 for arboviruses (flaviviruses and alphaviruses) and coronaviruses respectively. Supplemental Material provides a more detailed description of the assay.

### NOD2 silencing

Cells were seeded in 96-well plates (Greiner) overnight before transfection with 20 nM of NOD2 DsiRNA hs.Ri.NOD2.13.2 from Integrated DNA Technologies (IDT, Coralville, IA) using 0.3 μg/well RNAiMax (Invitrogen, Waltham, MA). The non-targeting IDT control DsiRNA was used as negative control for transfection. Twenty-four hours later, cells were infected with ZIKV using MOI of 0.05. At 24 and 48-hours post-infection, culture supernatants were collected for plaque assay. Total RNA isolated from cells at 48-hours post-infection was subjected to qRT-PCR to determine levels of viral genomic RNA and ISGs.

### Measurement of cell viability

Cell viability assays in response to drug or DMSO treatment were performed using CellTiter-Glo Luminescent Cell Viability kit (Promega) in cells grown in 96-well plates (Greiner) as described in the Supplemental Material.

### *In vitro* and *ex vivo* drug assays

After drug or DMSO treatment, viral replication and titers were determined by qRT-PCR on total RNA extracted from cells or tissues and plaque assay of culture supernatants respectively 12 to 72-hours post-infection. Cells seeded into 96-well plates (Greiner) were infected with ZIKV, DENV-2, MAYV or SARS-CoV-2 (MOI=0.05-5) followed by treatment with 5, 10, 20 and 40 μM GSK717 (14) (Sigma-Aldrich) or DMSO.

Fetal brain tissue explants were treated after ZIKV infection with GSK717 (20-40 μM) or DMSO for 3 days. Viral titer determination in culture supernatants daily and viral genome quantification in tissues at 72 hours post-infection were performed.

A549 or ACE2-SK-N-SH cells on coverslips in 12-well plates (Greiner) were infected (MOI of 1.0) with arboviruses (ZIKV, DENV-2 or MAYV) or SARS-CoV2 respectively and then processed for indirect immunofluorescence. Arbovirus co-infection assays and time-of-addition assays were conducted in A549 cells while SARS-CoV-2 time-of-addition assays were performed in ACE2-SK-N-SH using an MOI of 0.1. Viral genome quantification and viral titer determination were performed in the co-infection and time-of-addition assays, respectively.

A549 cells infected with arboviruses or ACE2-SK-N-SH cells infected with SARS-CoV-2 (MOI=0.05-5) were treated with the RIPK2 inhibitor GSK583 (28) (Sigma-Aldrich) for 24 hours. Cell supernatants and total cellular RNA were collected for determining viral titers and viral RNA respectively. Please, see Supplemental Material for additional information about the drug assays.

### Immunofluorescence staining and cell imaging

Infected cells grown on coverslips were fixed with 4% paraformaldehyde and permeabilized/blocked with a Triton-X100 (0.2%)/BSA (3%) solution and then incubated with mouse anti-Flavivirus Group Antigen 4G2 (Millipore, Burlington, MA), mouse anti-alphavirus capsid (kindly provided by Dr. Andres Merits at University of Tartu), or mouse anti-spike SARS-CoV/SARS-CoV-2 (GenTex, Irvine, CA) at room temperature for 1.5 hour, washed and then incubated with Alexa Fluor secondary antibodies against mouse and DAPI for 1 hour at room temperature. Antibodies were diluted in Blocking buffer. PBS containing 0.3% BSA was used for wash steps. Samples were examined using an Olympus 1×81 spinning disk confocal microscope (Tokyo, Japan) or Cytation 5 Cell Imaging Multi-Mode Reader instrument (Biotek, Winooski, VT). Images were analyzed using Volocity or Gen5 software. More experimental details are provided in the Supplemental Material.

### Statistical analyses

A paired Student’s t-test was used for pair-wise statistical comparison. The standard error of the mean is shown in all bar graphs. GraphPad Prism software 5.0 (GraphPad Software Inc., La Jolla, CA) was used in all statistical analyses.

## ACKNOWLEDGEMENTS

This work was supported by grants from the Canadian Institutes of Health Research (grants PJT-148699, OV3-172302, PJT-162417) and the Li Ka Shing Institute of Virology to T. C. H. The funders had no role in study design, data collection and interpretation, or the decision to submit the work for publication.

We wish to thank Dr. Anil Kumar, Dr. Joaquin Lopez Orozco, Dr. Adriana Airo, Dr. Zaikun Xu, Cheung Pang Wong, and Ray Ishida in the Hobman lab for helpful discussions and support. Confocal imaging was performed at the Cell Imaging Centre core in the Faculty of Medicine & Dentistry of the University of Alberta. We thank Dr. Darryl Falzarano (Vaccine and Infectious Disease Organization-International Vaccine Centre, University of Saskatchewan, Canada) for providing the Canadian SARS-CoV-2 strain for this work, Dr. David Safronetz (Public Health Agency of Canada, Canada) for providing the ZIKV strain, Dr. Bart Bartenschlager (Heidelberg University Hospital, Germany) for donating the DENV-2 clone, Dr. Eva Gönczöl (The Wistar Institute, USA) for providing the HEL-18 cells, and Dr. Andres Merits (University of Tartu, Estonia) for kindly donating the anti-alphavirus capsid antibody. Furthermore, we thank Eileen Reklow and Valeria Mancinelli for excellent technical support.

We declare no competing financial interests.

