## Supplemental Material for "Nodosome inhibition as a novel broad-spectrum antiviral strategy against arboviruses and SARS-CoV-2"

**Cells and viruses.** The prototype Zika virus (ZIKV) strain isolated in Puerto Rico (PRVABC-59) in 2015 (1) was kindly provided by the National Microbiology Laboratory of Canada (Winnipeg, Canada). Dengue virus-2 (DENV-2, 16681 strain) clone was provided by Dr. Bart Bartenschlager at the Heidelberg University Hospital (Heidelberg, Germany). Mayaro virus (MAYV, TRVL 15537) strain was obtained from ATCC. Arboviruses were propagated in *Aedes albopictus* C6/36 cells grown in MEM (Thermo Fisher Scientific) supplemented with 10% fetal bovine serum (Thermo Fisher Scientific), L-glutamine, Penicillin-Streptomycin and MEM non-essential amino acids at 32°C.

Arbovirus stocks were prepared after inoculating C6/36 cells using a multiplicity of infection (MOI) of 0.2 and then harvesting cell culture media between 24 to 96 hours post-infection. Virus-containing media were clarified by centrifugation at 3200 x g for 10 minutes. Human fetal astrocytes (HFAs) were prepared as previously described (2). Briefly, human fetal brain tissues were obtained from 15 to 19-week aborted fetuses. After isolation of astrocytes, cells were split, and all experiments were conducted with cells between the fifth and seventh passages. HFAs were grown in MEM supplemented with 10% fetal bovine serum, 15 mM HEPES (Thermo Fisher

Scientific) L-glutamine, MEM non-essential amino acids, sodium pyruvate, and 1g/mL glucose. Vero cells (ATCC, CCL-81) were maintained in DMEM supplemented with 10% fetal bovine serum, 15 mM HEPES, L-glutamine and Penicillin-Streptomycin.

Severe acute respiratory syndrome coronavirus 2 (SARS-CoV-2/CANADA/VIDO 01/2020) was kindly provided by Dr. Darryl Falzarano (Vaccine and Infectious Disease Organization-International Vaccine Centre, University of Saskatchewan, Canada). SARS-CoV-2 strain was propagated in Vero-E6 cells (ATCC) grown in DMEM supplemented with 10% fetal bovine serum, 15 mM HEPES, L-glutamine and Penicillin-Streptomycin.

A549, Huh7, and U251 (ATCC) and ACE2-hyperexpressing SK-N-SH cells were maintained in DMEM while HEL-18 human primary embryonic pulmonary fibroblasts (3) and Calu-3 (ATCC) were maintained in RPMI 1640 (Thermo Fisher Scientific) medium and MEM respectively supplemented with 10% fetal bovine serum, 15 mM HEPES, L-glutamine and Penicillin-Streptomycin.

**Generation of ACE2-SK-N-SH cells.** SK-N-SH cells stably expressing ACE2 were constructed using lentivirus transduction (4, 5). The coding region of the human ACE2 gene from the plasmid hACE2 (Addgene plasmid #1786) (6) was subcloned into the lentiviral vector pLenti-puro (Addgene plasmid #39481) (7) to generate pLenti-ACE2. Next, lentiviral pseudoparticles were harvested from the media of HEK293T cells (ATCC) transfected with viral packaging vectors (5) and pLenti-ACE2 using Lipofectamine 3000 (Invitrogen). SK-N-SH cells (ATCC) were transduced with the lentiviral pseudoparticles and stably transduced cells were selected using 10 μg/mL puromycin in DMEM containing 10% FBS. Expression of ACE2 protein was confirmed by western blotting using anti-ACE2 antibodies (R & D Systems).

**Viral infections of cells.** Cells were seeded in 96-wells plates (Greiner) at 1-2 x $10^4$ cells per well. The next day, cells were rinsed once with PBS and viruses at an MOI of 0.05 to 5 were added to the wells. Cells were then incubated for 1 (SARS-CoV-2), 2 (arboviruses) or 3 (arboviral co-infections) hours at 37°C using fresh media supplemented with 3% fetal bovine serum. Next, the inoculum was removed, and the cells were washed twice with PBS. Complete culture medium was added to each well, and cells were incubated at 37°C and 5% $CO_2$. Mock-infected cells were incubated with the culture supernatant from uninfected cells.

**Fetal brain explant cultures and infections.** Human brain tissue from two 15-19-week aborted fetuses was obtained (after written consent) under protocol 1420 of the University of Alberta Human Research Ethics Board. After delivery to the laboratory in ice-cold PBS, the fresh tissue was placed in a 100 mM Petri dish and then dissected with sterile scalpel and forceps into approximately 5 mm x 5 cm x 1 mm blocks (2). The small tissue blocks were immediately immersed in 24-well plates containing the same media used to culture HFAs (see above).

Brain explant tissue was infected overnight with $10^6$ PFU/mL of PRVABC-59 ZIKV strain. The next day, explants were washed once with PBS and then fresh media was added. The explant cultures were maintained in a humidified 37°C incubator containing 5% $CO_2$ for up to 5 days under different experimental treatments.

**Tissue and cellular RNA purification, cDNA synthesis, and qRT-PCR.** Total RNA was extracted from cultured cells or fetal brain tissue using NucleoSpin RNA (Macherey-Nagel GmbH & Co) kits. Samples were then treated with RNase-free DNase (Macherey-Nagel GmbH & Co) before a portion (0.5-1 µg total RNA) was subjected to reverse transcription using ImProm-II Reverse Transcriptase (Promega). Cellular transcripts and viral RNA were quantitated by qRT-

PCR using PerfeCTa SYBR Green SuperMix (Quanta BioSciences) in a CFX96 Touch Real-Time PCR Detection System instrument (Bio-Rad) under the following cycling conditions: 40 cycles of 94°C for 30 s, 55°C for 60 s, and 68°C for 20 s. Gene expression (fold change) was calculated using the $2^{(-\Delta\Delta CT)}$ method with human β-actin mRNA transcript as the internal control.

The following forward and reverse primer pairs were used for PCR:

| **Primer name** | **Sequence** |
|---|---|
| *β-actin* | 5'-GGATCAGCAAGCAGGAGTATG-3'<br>5'-GCATTTGCGGTGGACGAT-3' |
| *NOD2* (nucleotide-binding oligomerization domain-containing protein 2) | 5'-TCTCTGTGCGGACTCTACTC-3'<br>5'-ATCCGTGAACCTGAACTTG-3' |
| *RSAD2* (radical SAM domain-containing 2 or viperin) | 5'-CTTTTGCTGGGAAGCTCTTG-3'<br>5'-CAGCTGCTGCTTTCTCCTCT-3' |
| *GSDMD* (gasdermin D) | 5'-GTGTGTCAACCTGTCTATCAAGG-3'<br>5'-CATGGCATCGTAGAAGTGGAAG-3' |
| *Casp1* (caspase 1) | 5'-TCACTGCTTCGGACATGACTACA-3'<br>5'-GGAACGTGCTGTCAGAGGTCTT-3' |
| *GBP5* (guanylate binding protein 5) | 5'-TCCTCGGATTATTGCTCGGC-3'<br>5'- CCTTTGCGCTTCAGCCTTTT-3' |
| *NLRC5* (NOD-like receptor family CARD domain containing 5) | 5'-TGGGAAGACACTCAGGCTAA-3'<br>5'-ATCATCGTCCTCACAGAGGTT-3' |
| *IL-18* (Interleukin-18) | 5'- GACTGTAGAGATAATGCAC-3' |

| | |
|---|---|
| | 5’-CTTCGTTTTGAACAGTGAAC-3’ |
| *NLRP1* (NLR Family Pyrin Domain Containing 1) | 5’-ACCATGGTAGTCCTGTTCAG-3’<br>5’-GCGAGTTTCCACTTAGGTC-3’ |
| *ASC* (apoptosis-associated speck like protein containing a caspase recruitment domain) | 5’-GCACTTTATAGACCAGCACCG-3’<br>5’-CTGAAGAGCTTCCGCATCTTG-3’ |
| *IL-1 β* (Interleukin-1 β) | 5’-AACCTCTTCGAGGCACAAGG-3’<br>5’-GTCCTGGAAGGAGCACTTCAT-3’ |
| *EIF2AK2* (eukaryotic translation initiation factor 2-alpha kinase 2) | 5’-ACCTCAGTGAAATCTGACTACC-3’<br>5’-CAGATGATGATTCAGAAGCG -3’ |
| *IFI-16* (gamma-interferon-inducible protein 16) | 5’- ACAAACCCGAGAAACAATGACC-3’<br>5’- GCATCTGAGGAGTCCGAAGA-3’ |
| *NLRC4* (NOD-like receptor family CARD domain containing 4) | 5’-CTGAGCAGCCTGTTGAAAC-3’<br>5’-CATCCATCACTGCTCACAC-3’ |
| *NLRP3* (NLR family pyrin domain containing 3) | 5’-CCAAGAATCCACAGTGTAACC-3’<br>5’-CTTCACAGAACATCATGACCC-3’ |
| *OAS-1* (2'-5'-oligoadenylate synthetase 1) | 5’- TTCTTAAAGCATGGGTAATTC-3’<br>5’- GAAGGCAGCTCACGAAAC-3’ |
| *MX2* (Myxovirus resistance protein 2 or interferon-induced GTP-binding protein MX2) | 5’-CAGCCACCACCAGGAAACA-3’<br>5’-TTCTGCTCGTACTGGCTGTACAG-3’ |
| ZIKV (8) | 5’-CCTTGGATTCTTGAACGAGGA-3’<br>5’-AGAGCTTCATTCTCCAGATCAA-3’ |

| | |
|---|---|
| DENV (9) | 5’-TTGAGTAAACTGTGCAGCCTGTAGCTC-3’<br><br>5’-GGGTCTCCTCTAACCTCTAGTCCT-3’ |
| MAYV (10) | 5’-AAGCTCTTCCTCTGCATTGC-3’<br><br>5’-TGCTGGAAACGCTCTCTGTA-3’ |
| SARS-CoV-2 (11) | 5’-CAATGGTTTAACAGGCACAGG-3’<br><br>5’-CTCAAGTGTCTGTGGATCACG-3’ |

**Viral titer assay.** Flaviviruses and alphaviruses were serially diluted (10-fold dilutions) and monolayers of Vero CCL-81 cells at 37 °C were infected for 2 hours. For the SARS-CoV-2 plaque assay, Vero-E6 cells were infected for 1 hour in a biosafety level 3 laboratory of the University of Alberta. The monolayers were overlaid with a mixture of MEM (Thermo Fisher Scientific) and 0.75-1.5% carboxymethylcellulose (Sigma-Aldrich) following infection. The cells were maintained at 37 °C for 2-7 days, depending on the virus (2 days for MAYV, 3 days for SARS-CoV-2, 4 days for ZIKV, and 7 days for DENV-2) for plaque development. Cells were fixed with 10% formaldehyde and stained with 1% crystal violet in 20% ethanol after which plaques were counted.

**Cytotoxicity assays.** The CellTiter-Glo Luminescent Cell Viability Assay (Promega) was used for quantitation of ATP in cultured cells. Cells lysates were assayed after mixing 100 µl of complete media with 100 µl of reconstituted CellTiter-Glo Reagent (buffer plus substrate) following the manufacturer’s instructions. Samples were mixed by shaking the plates after which luminescence was recorded with a GloMax Explorer Model GM3510 (Promega) 10 min after adding the reagent.

**Antiviral drug assays.** Unless otherwise indicated, cells were cultured in 96-well plates (Greiner) and viral replication and titers were determined by qRT-PCR on total RNA extracted from cells and plaque assay of cell supernatants respectively. Cells seeded into 96-well plates (Greiner) at 1-2 x $10^4$ cells per well were infected the next day with ZIKV, DENV-2, MAYV or SARS-CoV-2 (MOI=0.05-5) followed by treatment with 5, 10, 20 and 40 µM of GSK717 (12) (Sigma-Aldrich) or an equal volume of DMSO. Viral replication and titers were determined 24 to 72-hours post-infection.

HFAs (5 x $10^4$ cells per well) were grown in 24-well plates (Greiner) and after ZIKV infection (MOI=0.05-5), GSK717 (5-40 µM) was added to carry out a 3-day kinetics of viral titers. ZIKV-infected fetal brain tissue explants were treated with GSK717 (20 and 40 µM) or DMSO for 3 days. Viral titer determination in culture supernatants by plaque assay daily and viral genome quantification in tissues by qRT-PCR at 72 hours post-infection were performed.

For some drug assays, an MOI of 1.0 was used to infect A549 cells on coverslips with ZIKV, DENV-2 or MAYV at 1 x $10^5$ cells per well in 12-well plates (Greiner) for indirect immunofluorescence of viral antigens. For the drug assays in co-infected A549 cells (DENV-2 and ZIKV or DENV-2 and MAYV), an MOI of 0.1 was used. For time-of-addition drug assays, nodosome inhibitors or DMSO were added either immediately or 12 or 24-hours after the virus absorption step. Cell supernatants were collected for viral titer determination at 24, 36, and 48-hours after addition of the compounds.

After infecting A549 cells with ZIKV, DENV-2 or MAYV (MOI=0.05-5), cells were treated with the RIPK2 inhibitor GSK583 (13) (Sigma-Aldrich) or DMSO as control for 24 hours. Furthermore, a time-of-addition assay, as described above, was performed. Cell supernatants, cells on coverslips and total cellular RNA were collected for determining viral titers by plaque assay, percentage of infected cells by indirect immunofluorescence and viral replication by qRT-PCR respectively.

GSK717 or GSK583 were used to treat SARS-CoV-2 (MOI=0.05-5) infected ACE2-SK-N-SH cells seeded into 96-well or 12-well-plates, followed by viral RNA quantification, viral titer determination or indirect immunofluorescence as described above for the arbovirus inhibition assays. Calu-3 and Huh7 cells infected with SARS-CoV-2 (MOI=0.1) were also treated with GSK583 for 24 hours before collecting the cell supernatants for viral titer determination.

As a positive control of the flaviviral inhibition in infected A549 cells (MOI=0.5-5), we used the anti-flavivirus nucleoside analog NITD008 (14) (Sigma-Aldrich) at 0.75-3 µM or DMSO alone. In the SARS-CoV-2 inhibition assays, remdesivir (15) (MedKoo Biosciences, Inc.) in infected Vero E6 cells (MOI=0.1) was used as positive control at 0.1-10 µM or DMSO as vehicle.

**Immunostaining and imaging.** Cells on coverslips were processed by fixing for 15 min at room temperature with 4% paraformaldehyde in PBS. Cells were washed three times in PBS and then permeabilized/blocked in Blocking buffer with 0.2% Triton-X100 and 3% BSA in PBS for 1 hour at room temperature followed by washing with PBS containing 0.3% BSA. Incubations with primary antibodies diluted 1:500 (mouse anti-Flavivirus Group Antigen 4G2, Millipore), 1:200 (mouse monoclonal anti-chikungunya capsid kindly donated by Dr. Andres Merits at University of Tartu, Estonia)(16), or 1:250 (mouse monoclonal anti-spike SARS-CoV/SARS-CoV-2, GenTex) in Blocking buffer were carried out at room temperature for 1.5 hour followed by three washes in Washing buffer (0.02% Triton-X100 with 0.3% BSA and PBS).

Samples were then incubated with secondary antibodies (1:1000) in Blocking buffer containing 1 µg/mL of DAPI for 1 hour at room temperature followed by three washes in Washing buffer. Secondary antibodies (Invitrogen) were Alexa Fluor 488 anti-mouse. Confocal images of cells on coverslips were acquired using an Olympus 1x81 spinning disk confocal microscope (Tokyo, Japan) and images were analyzed using Volocity 6.2.1 software. A set of confocal imaging was acquired using a Cytation 5 Cell Imaging Multi-Mode Reader and analyzed using Gen5 software (Biotek). Total and antigen-positive cells were counted in 10 image fields per well, and the percentage of infected cells was obtained.

**Supplemental Figure legends**

**Fig S1. Inflammasome induction by human recombinant IFN-α and NOD2 silencing in HFAs.** HFAs were treated with human recombinant IFN-α for 4, 8 and 12 hours after which relative expression of inflammasome genes gasdermin D (*GSDMD*) (**A**), caspase 1 (*Casp1*) (**B**), guanylate binding protein 5 (*GBP5*) (**C**) and NOD-like receptor family CARD domain containing 5 (*NLRC5*) (**D**) were determined. (**E**) HFAs were transfected with NOD2-specific or non-silencing siRNAs for 24-hours followed by ZIKV infection (MOI=0.05). Total cellular RNA was collected at 48 hours post-infection for *NOD2* gene quantitation by qRT-PCR. Relative levels of *NOD2* is shown. (**F**) Cellular ATP levels in uninfected HFAs were determined using the CellTiter-Glo assay kit after *NOD2* silencing for 72 hours. Values are expressed as the mean of three independent experiments. Error bars represent standard errors of the mean. *$P < 0.05$, **$P < 0.01$, and ***$P < 0.001$, by the Student *t* test.

**Fig S2. NOD2 blocking drug GSK717 displays anti-ZIKV activity in different cell types.** Human primary embryonic pulmonary fibroblasts (HEL-18) were treated with GSK717 or DMSO as control for 72 hours after ZIKV infection with MOI of 0.05 and 5. ZIKV titers are shown as relative fold with MOI of 0.05 (**A)** and 5 (**B**) at 48-72 hours post-infection and as PFU/mL with MOI of 0.05 (**C**) at 48 hours post-infection. ZIKV titers as relative fold in A549 (**D**), U251 (**E**) and Huh7 (**F**) cells after 48 hours of infection (MOI=0.05-5) with and without GSK717 treatment are shown. (**G**) Viral titers as relative fold in A549 cells infected with ZIKV (MOI=0.1) for 0, 12 or 24 hours followed by GSK717 drug or DMSO addition for 48, 36 or 24 hours respectively are shown. Values are expressed as the mean of three independent experiments. Error bars represent standard errors of the mean. *$P < 0.05$, **$P < 0.01$, and ***$P < 0.001$, by the Student *t* test.

**Fig S3. Anti-NOD2 drug is not cytotoxic in human primary cells and cell lines.** (**A**) Cellular ATP levels, measured by the CellTiter-Glo assay kit, of uninfected HFAs treated with the GSK717 or DMSO for 72 hours are shown. Cellular ATP measurements in the A549 (**B**), U251 (**C**), Huh7 (**D**) and ACE2-SK-N-SH (**E**) cell lines after 48 hours of GSK717 or DMSO treatment are presented. Values are expressed as the mean of three independent experiments. Error bars represent standard errors of the mean.

**Fig S4. GSK717 blocks the spread of DENV-2.** (**A**) Representative confocal imaging (20X) of DENV-2 infected cells treated with GSK717. A549 cells were infected with DENV-2 (MOI=1) followed by treatment with DMSO or GSK717 at 20 or 40 µM for 48 hours before fixation and immunostaining. DENV-infected cells were detected using mouse monoclonal antibody (4G2) to envelope protein and Alexa Fluor 488 donkey anti-mouse. Nuclei were stained with DAPI. Images were acquired using a spinning disk confocal microscope equipped with Volocity 6.2.1 software. (**B**) Infected cells were counted in 10 different fields for GSK717 and DMSO treated samples. Values are expressed as the mean of three independent experiments. Error bars represent standard errors of the mean. **$P < 0.01$, by the Student *t* test.

**Fig S5. RIPK2 inhibitor GSK583 suppresses arboviral replication at sub-cytotoxic concentrations.** A549 cells were infected with MAYV (MOI=0.1) and treated with RIPK2 inhibitor GSK583 (30 µM) or DMSO. Relative fold of titers (**A**) and viral genome (**B**) at 12- and 24-hours post-infection is shown. (**C**) A549 cells infected separately with arboviruses at low (0.05) and high MOI (5) were treated with GSK583 (30 µM) or DMSO for 24 hours before supernatant collection for plaque assays. Viral titers as relative fold is shown. (**D**) Cellular ATP measurements by the CellTiter-Glo assay kit in uninfected A549 after 24 hours of GSK583 or DMSO treatment are shown. Values are expressed as the mean of three independent experiments. Error bars represent standard errors of the mean. $*P < 0.05$, $**P < 0.01$, and $***P < 0.001$ by the Student *t* test.

**Fig S6. RIPK2 inhibitor GSK583 blocks the spread of arboviruses**. (**A**) Representative confocal imaging (20X) of GSK583 effect on MAYV-infected A549 cells. Cells were infected with MAYV, ZIKV or DENV (MOI=1) and then treated with GSK583 (15 or 30 µM) for 24 hours before processing for indirect immunofluorescence. Infected cells were identified using mouse anti-alphavirus capsid (MAYV) or anti-flavivirus envelope 4G2 (ZIKV and DENV-2) and Alexa Fluor 488 donkey anti-mouse. Nuclei were stained with DAPI. Images of MAYV-infected cells were acquired using a Cytation 5 Cell Imaging Multi-Mode Reader and analyzed using Gen5 software (Biotek) while images of flavivirus infected cells (ZIKV and DENV-2) were acquired using a spinning disk confocal microscope with Volocity 6.2.1 software. Infected cells were determined by counting within 10 different fields for each sample of MAYV (**B**), ZIKV (**C**) and DENV-2 (**D**). Values are expressed as the mean of three independent experiments. Error bars represent standard errors of the mean. $*P < 0.05$, $**P < 0.01$, and $***P < 0.001$, by the Student *t* test.

**Fig S7. RIPK2 inhibitor blocks the infection by and spread of SARS-CoV-2 in ACE2-SK-N-SH.** (**A**) ACE2-SK-N-SH were infected with SARS-CoV-2 (MOI=0.1) followed by treatment with 30 µM of GSK583 or DMSO. Cell supernatants were collected at 12- and 24-hours post-infection for plaque assay. Viral titers as PFU/mL is shown. (**B**) The same drug concentration was also added at 0 and 24 hours after infection with SARS-CoV-2 (MOI=0.1) to determine titers (PFU/mL) in ACE2-SK-N-SH supernatants 24 hours post-treatment. (**C**) Representative confocal imaging

(20X) of ACE2-SK-N-SH infected with SARS-CoV-2 (MOI=1) and treated with GSK583 or DMSO for 24 hours. After fixation, coronavirus-infected cells were stained using mouse monoclonal antibody to SARS spike protein as primary antibody and secondary antibodies were Alexa Fluor 488 donkey anti-mouse. Nuclei were stained with DAPI. Confocal images were acquired using a spinning disk confocal microscope with Volocity 6.2.1 software. (**D**) Cells were counted in 10 different fields for quantitation of infected cells after GSK583 or DMSO treatment. Values are expressed as the mean of three independent experiments. Error bars represent standard errors of the mean. *$P < 0.05$, **$P < 0.01$, and ***$P < 0.001$, by the Student *t* test.

**Fig S8. GSK583 suppresses SARS-CoV-2 release at subtoxic concentrations.** Calu-3 (**A**) and Huh7 (**B**) cells were infected with SARS-CoV-2 at the MOI of 0.1 and treated with GSK583 or DMSO as control for 24 hours after which cell supernatants were collected for plaque assay. Viral titers as relative fold is shown. Cellular ATP measurements using the CellTiter-Glo assay kit are shown at the 24-hour time point after GSK583 or DMSO treatment in uninfected ACE2-SK-N-SH (**C**), Calu-3 (**D**), and Huh7 (**E**) cells. Values are expressed as the mean of three independent experiments. Error bars represent standard errors of the mean. *$P < 0.05$, **$P < 0.01$, and ***$P < 0.001$, by the Student *t* test.

11. Zhou P, Yang XL, Wang XG, Hu B, Zhang L, Zhang W, Si HR, Zhu Y, Li B, Huang CL, Chen HD, Chen J, Luo Y, Guo H, Jiang RD, Liu MQ, Chen Y, Shen XR, Wang X, Zheng

XS, Zhao K, Chen QJ, Deng F, Liu LL, Yan B, Zhan FX, Wang YY, Xiao GF, Shi ZL. 2020. A pneumonia outbreak associated with a new coronavirus of probable bat origin. Nature 579:270-273.

12. Rickard DJ, Sehon CA, Kasparcova V, Kallal LA, Zeng X, Montoute MN, Chordia T, Poore DD, Li H, Wu Z, Eidam PM, Haile PA, Yu J, Emery JG, Marquis RW, Gough PJ, Bertin J. 2013. Identification of benzimidazole diamides as selective inhibitors of the nucleotide-binding oligomerization domain 2 (NOD2) signaling pathway. PLoS One 8:e69619.

13. Haile PA, Votta BJ, Marquis RW, Bury MJ, Mehlmann JF, Singhaus R, Charnley AK, Lakdawala AS, Convery MA, Lipshutz DB, Desai BM, Swift B, Capriotti CA, Berger SB, Mahajan MK, Reilly MA, Rivera EJ, Sun HH, Nagilla R, Beal AM, Finger JN, Cook MN, King BW, Ouellette MT, Totoritis RD, Pierdomenico M, Negroni A, Stronati L, Cucchiara S, Ziółkowski B, Vossenkämper A, MacDonald TT, Gough PJ, Bertin J, Casillas LN. 2016. The Identification and Pharmacological Characterization of 6-(tert-Butylsulfonyl)-N-(5-fluoro-1H-indazol-3-yl)quinolin-4-amine (GSK583), a Highly Potent and Selective Inhibitor of RIP2 Kinase. J Med Chem 59:4867-80.

14. Yin Z, Chen YL, Schul W, Wang QY, Gu F, Duraiswamy J, Kondreddi RR, Niyomrattanakit P, Lakshminarayana SB, Goh A, Xu HY, Liu W, Liu B, Lim JY, Ng CY, Qing M, Lim CC, Yip A, Wang G, Chan WL, Tan HP, Lin K, Zhang B, Zou G, Bernard KA, Garrett C, Beltz K, Dong M, Weaver M, He H, Pichota A, Dartois V, Keller TH, Shi PY. 2009. An adenosine nucleoside inhibitor of dengue virus. Proc Natl Acad Sci U S A 106:20435-9.

15. Pruijssers AJ, George AS, Schäfer A, Leist SR, Gralinksi LE, Dinnon KH, Yount BL, Agostini ML, Stevens LJ, Chappell JD, Lu X, Hughes TM, Gully K, Martinez DR, Brown AJ, Graham RL, Perry JK, Du Pont V, Pitts J, Ma B, Babusis D, Murakami E, Feng JY, Bilello JP, Porter DP, Cihlar T, Baric RS, Denison MR, Sheahan TP. 2020. Remdesivir Inhibits SARS-CoV-2 in Human Lung Cells and Chimeric SARS-CoV Expressing the SARS-CoV-2 RNA Polymerase in Mice. Cell Rep 32:107940.
16. Goh LYH, Hobson-Peters J, Prow NA, Gardner J, Bielefeldt-Ohmann H, Suhrbier A, Hall RA. 2015. Monoclonal antibodies specific for the capsid protein of chikungunya virus suitable for multiple applications. J Gen Virol 96:507-512.

FIG S1

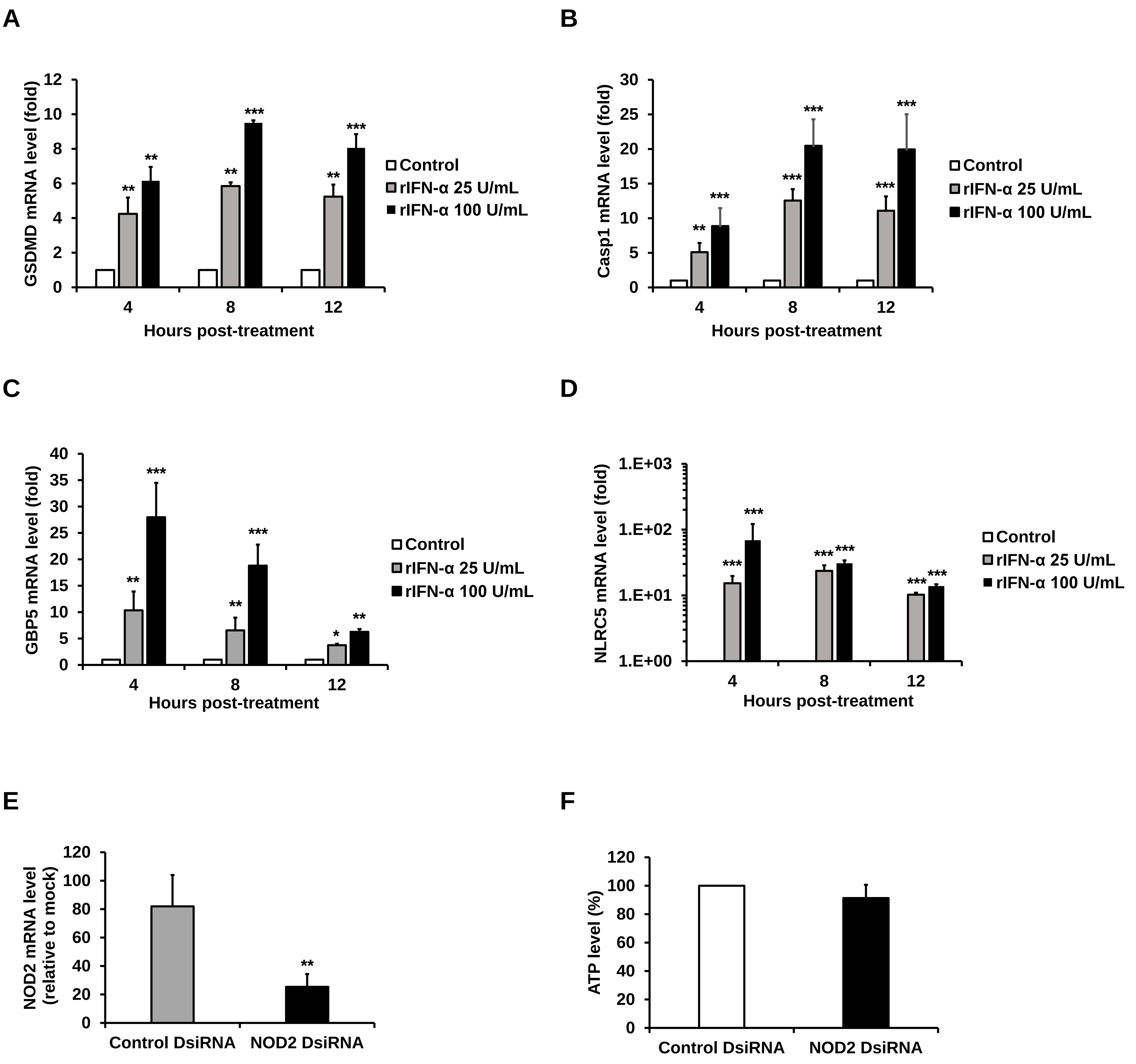

A
GSDMD mRNA level (fold)
0
2
4
6
8
10
12
Control
rIFN-α 25 U/mL
rIFN-α 100 U/mL
4
8
12
Hours post-treatment
B
Casp1 mRNA level (fold)
0
5
10
15
20
25
30
Control
rIFN-α 25 U/mL
rIFN-α 100 U/mL
4
8
12
Hours post-treatment
C
GBP5 mRNA level (fold)
0
5
10
15
20
25
30
35
40
Control
rIFN-α 25 U/mL
rIFN-α 100 U/mL
4
8
12
Hours post-treatment
D
NLRC5 mRNA level (fold)
1.E+00
1.E+01
1.E+02
1.E+03
Control
rIFN-α 25 U/mL
rIFN-α 100 U/mL
4
8
12
Hours post-treatment
E
NOD2 mRNA level (relative to mock)
0
20
40
60
80
100
120
Control DsiRNA
NOD2 DsiRNA
F
ATP level (%)
0
20
40
60
80
100
120
Control DsiRNA
NOD2 DsiRNA

FIG S2

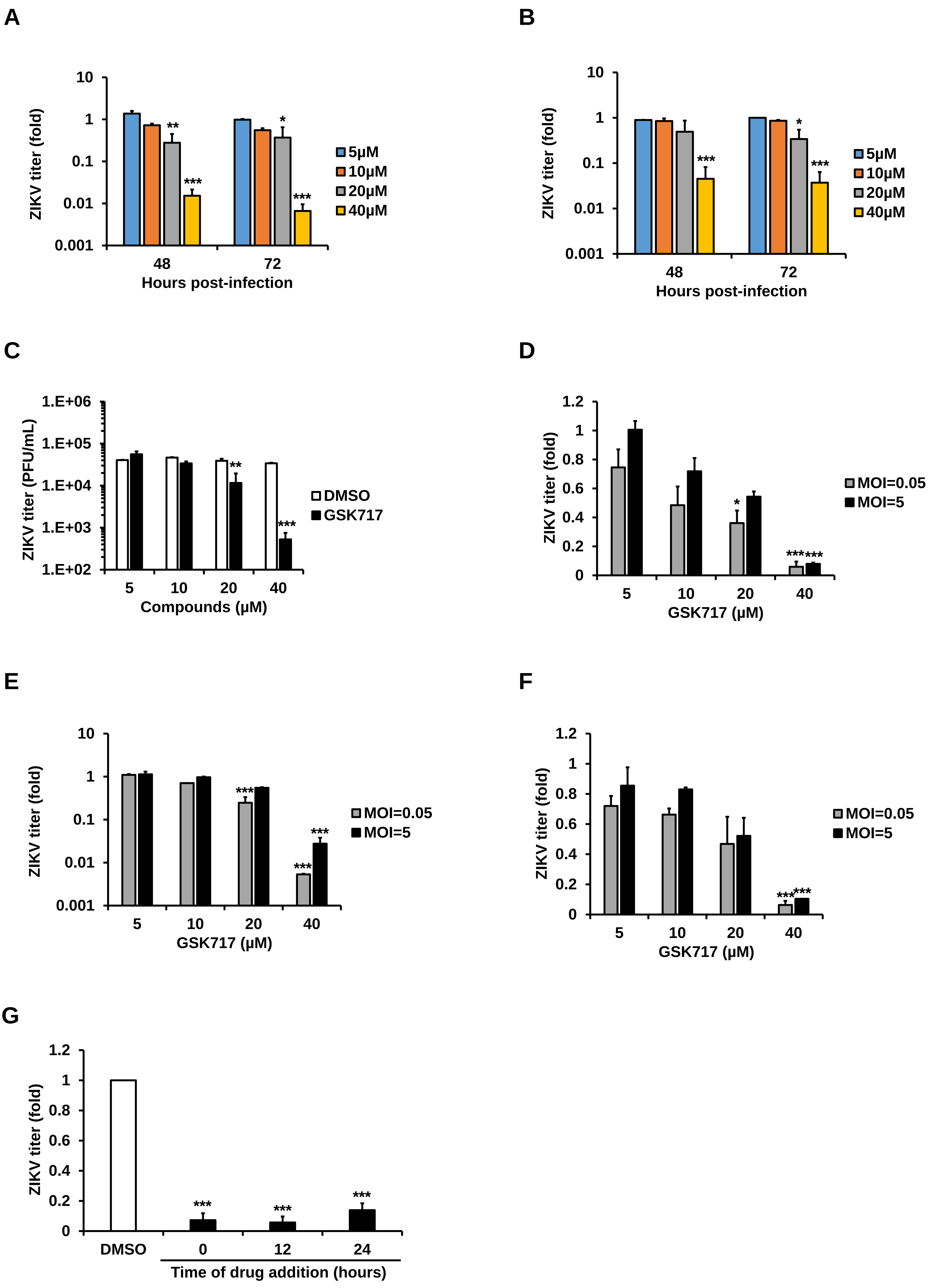
A
ZIKV titer (fold)
10
1
0.1
0.01
0.001
**
***
*
***
48
72
Hours post-infection
5μM
10μM
20μM
40μM
B
ZIKV titer (fold)
10
1
0.1
0.01
0.001
***
*
***
48
72
Hours post-infection
5μM
10μM
20μM
40μM
C
ZIKV titer (PFU/mL)
1.E+06
1.E+05
1.E+04
1.E+03
1.E+02
**
***
5
10
20
40
Compounds (μM)
DMSO
GSK717
D
ZIKV titer (fold)
1.2
1
0.8
0.6
0.4
0.2
0
*
******
5
10
20
40
GSK717 (μM)
MOI=0.05
MOI=5
E
ZIKV titer (fold)
10
1
0.1
0.01
0.001
***
***
***
5
10
20
40
GSK717 (μM)
MOI=0.05
MOI=5
F
ZIKV titer (fold)
1.2
1
0.8
0.6
0.4
0.2
0
******
5
10
20
40
GSK717 (μM)
MOI=0.05
MOI=5
G
ZIKV titer (fold)
1.2
1
0.8
0.6
0.4
0.2
0
***
***
***
DMSO
0
12
24
Time of drug addition (hours)

FIG S3

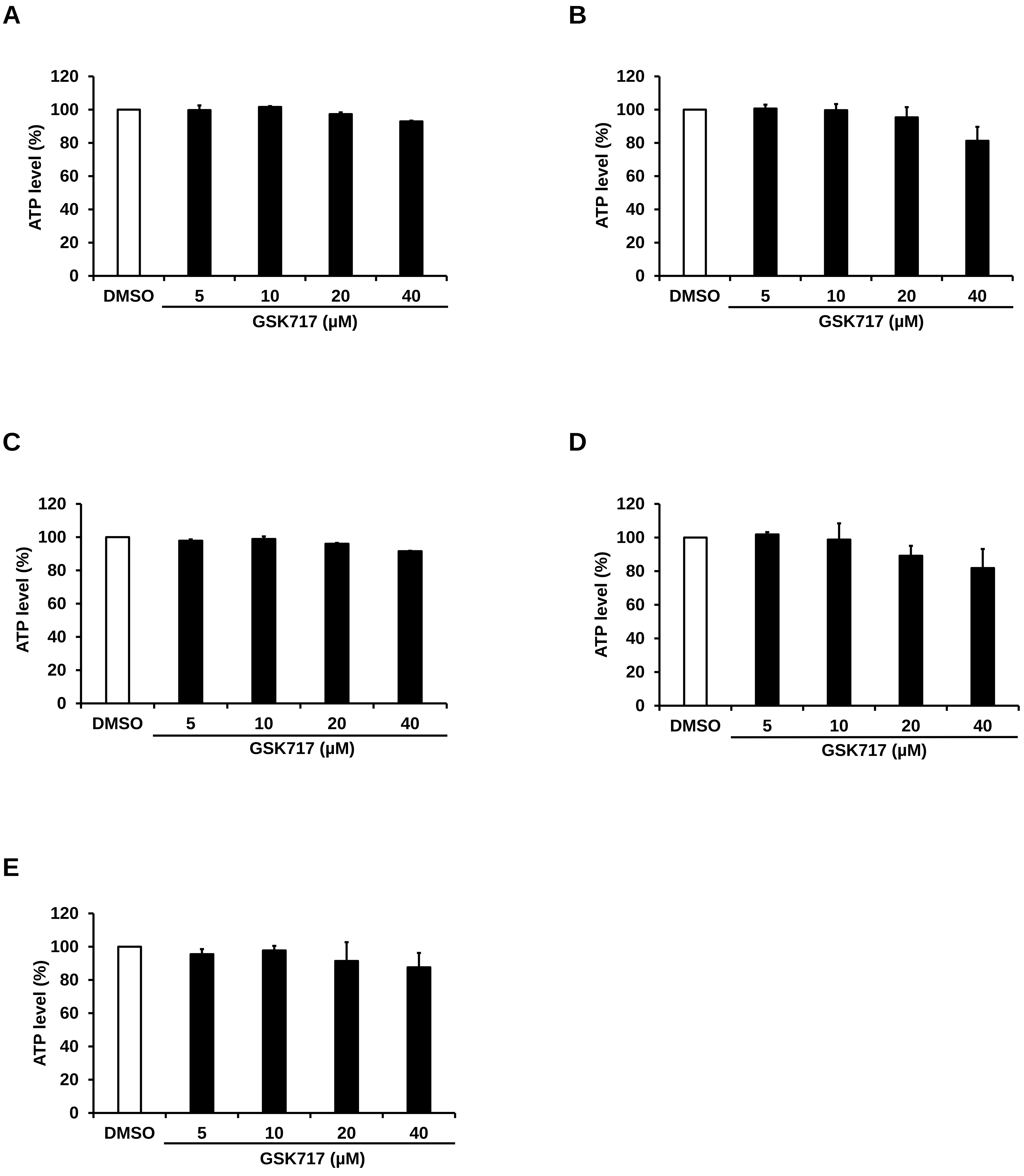
A
120
100
80
60
40
20
0
ATP level (%)
DMSO
5
10
20
40
GSK717 (μM)
B
120
100
80
60
40
20
0
ATP level (%)
DMSO
5
10
20
40
GSK717 (μM)
C
120
100
80
60
40
20
0
ATP level (%)
DMSO
5
10
20
40
GSK717 (μM)
D
120
100
80
60
40
20
0
ATP level (%)
DMSO
5
10
20
40
GSK717 (μM)
E
120
100
80
60
40
20
0
ATP level (%)
DMSO
5
10
20
40
GSK717 (μM)

FIG S4

A

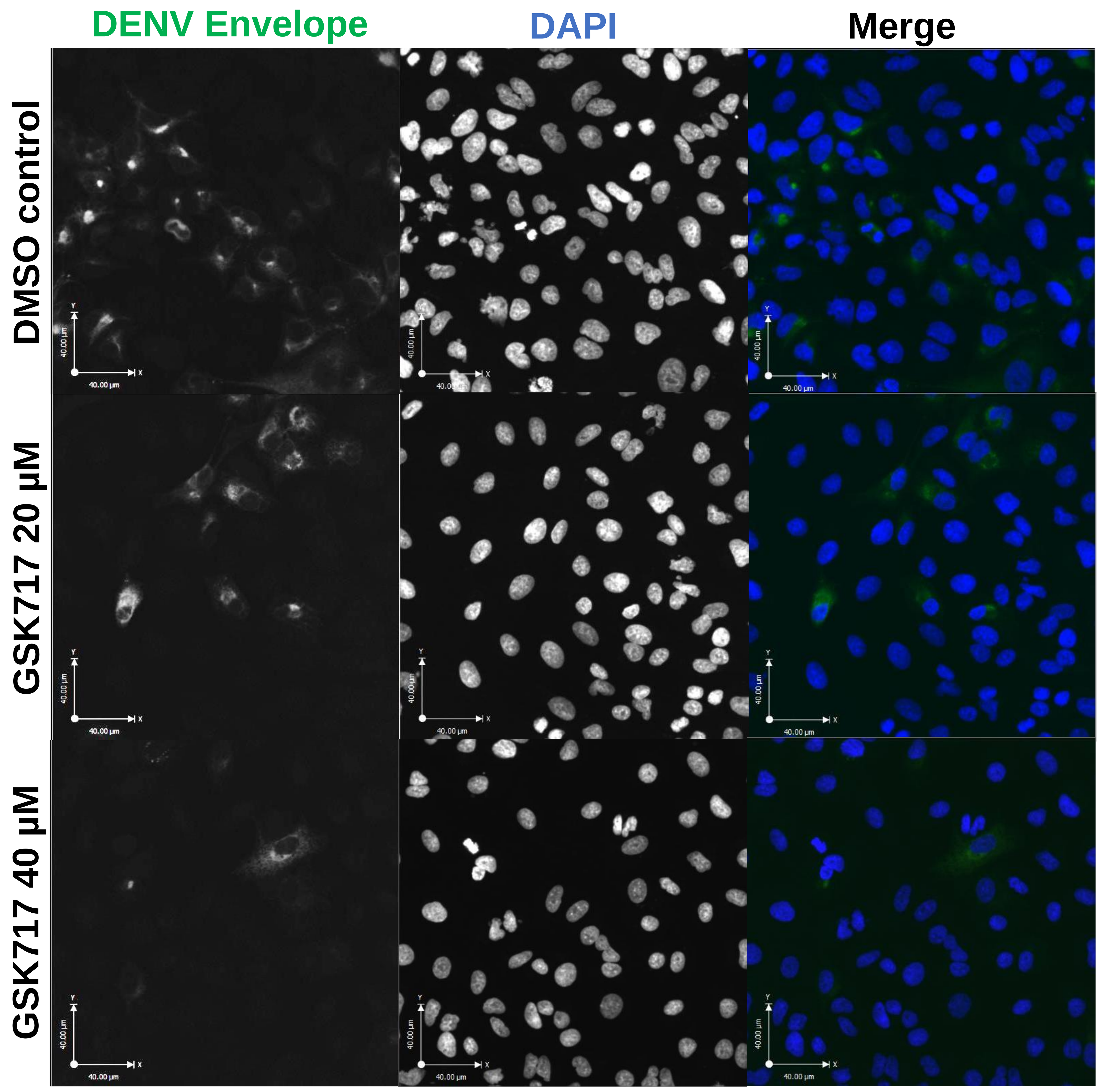
DENV Envelope
DAPI
Merge
DMSO control
GSK717 20 μM
GSK717 40 μM

B

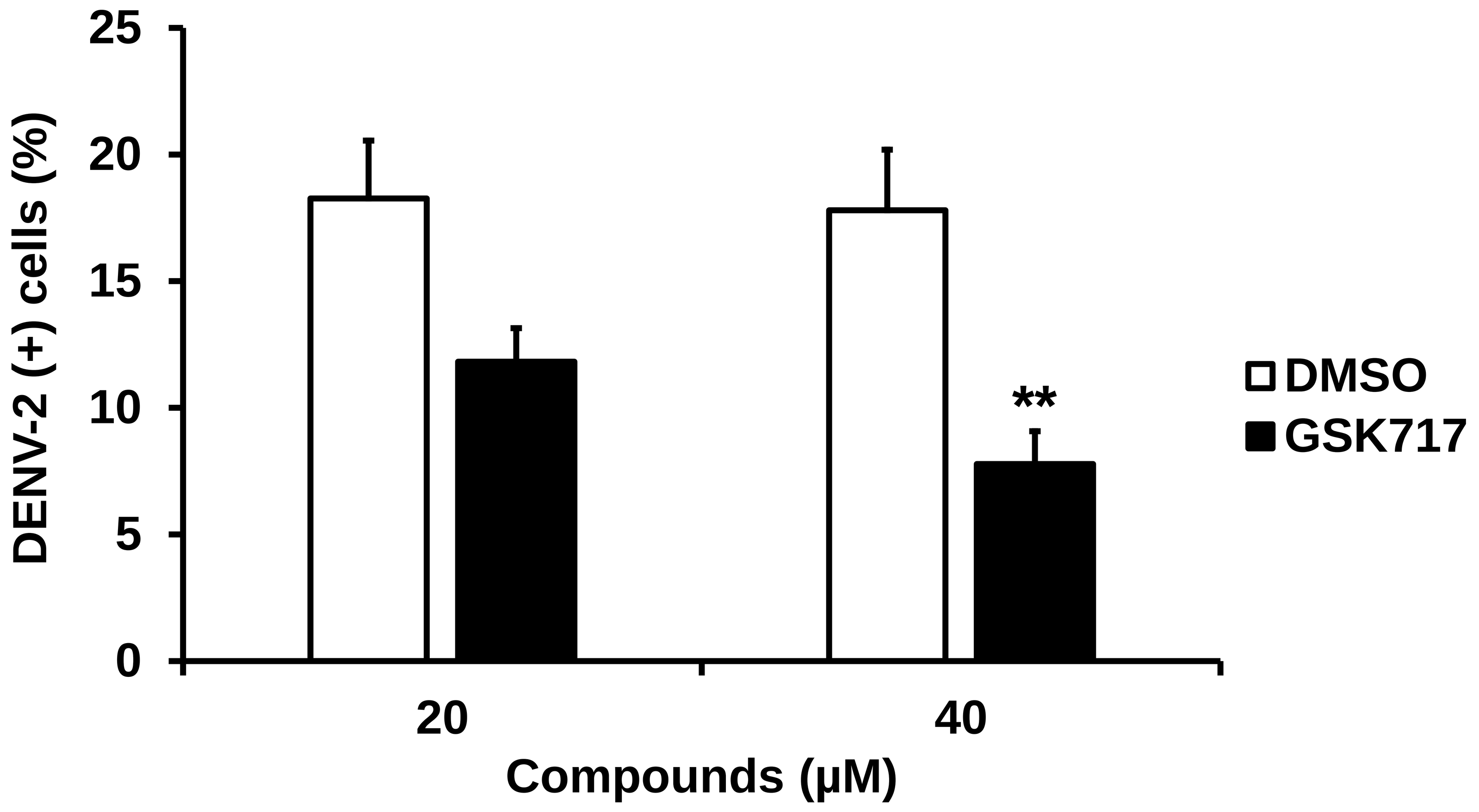
25
20
15
10
5
0
DENV-2 (+) cells (%)
**
DMSO
GSK717
20
40
Compounds (μM)

### FIG S5

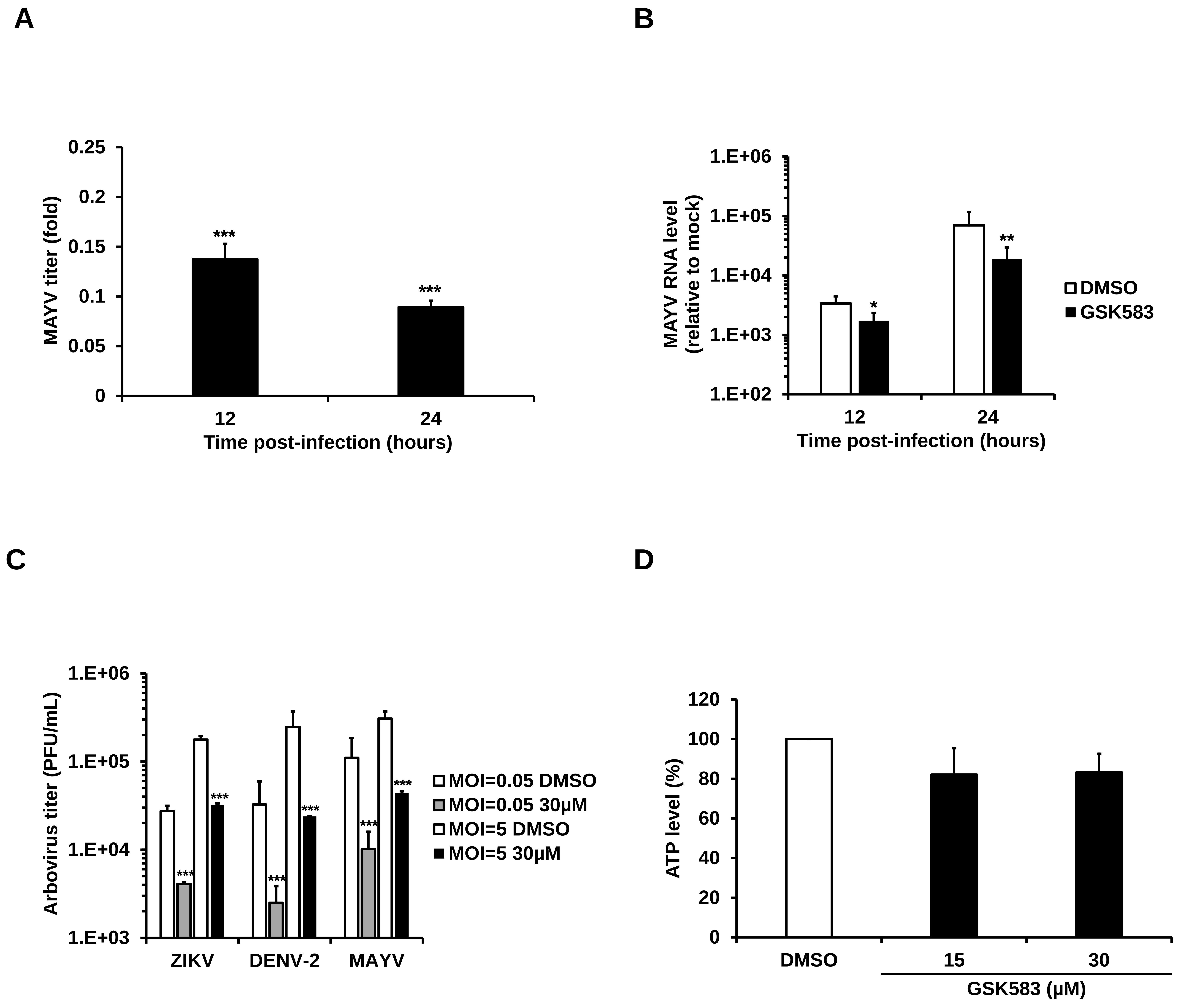
A
B
C
D
0.25
0.2
0.15
0.1
0.05
0
MAYV titer (fold)
***
***
12
24
Time post-infection (hours)
1.E+06
1.E+05
1.E+04
1.E+03
1.E+02
MAYV RNA level
(relative to mock)
*
**
DMSO
GSK583
12
24
Time post-infection (hours)
1.E+06
1.E+05
1.E+04
1.E+03
Arbovirus titer (PFU/mL)
***
ZIKV
DENV-2
MAYV
MOI=0.05 DMSO
MOI=0.05 30μM
MOI=5 DMSO
MOI=5 30μM
120
100
80
60
40
20
0
ATP level (%)
DMSO
15
30
GSK583 (μM)

FIG S6

A

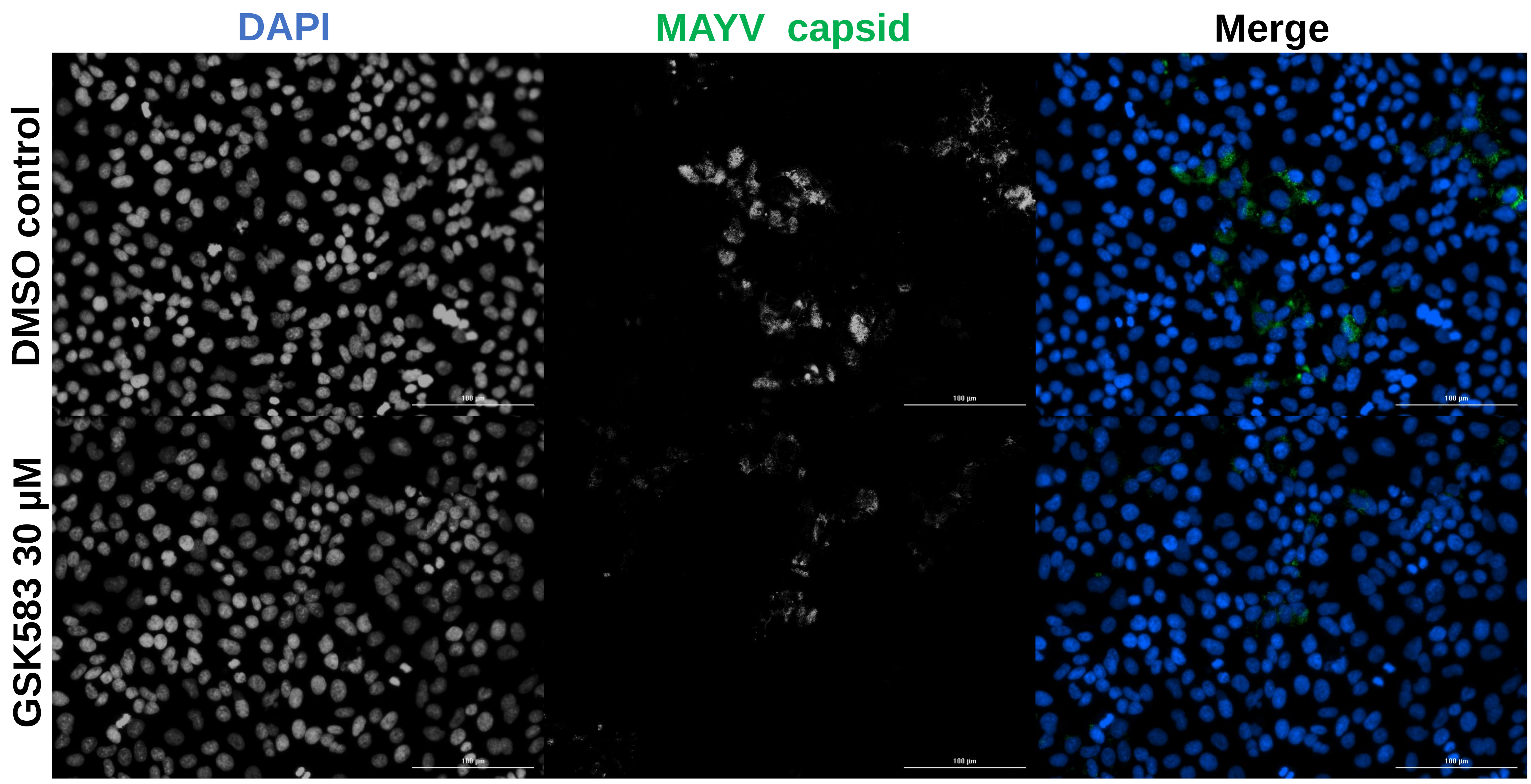
DAPI
MAYV capsid
Merge
DMSO control
GSK583 30 µM

B

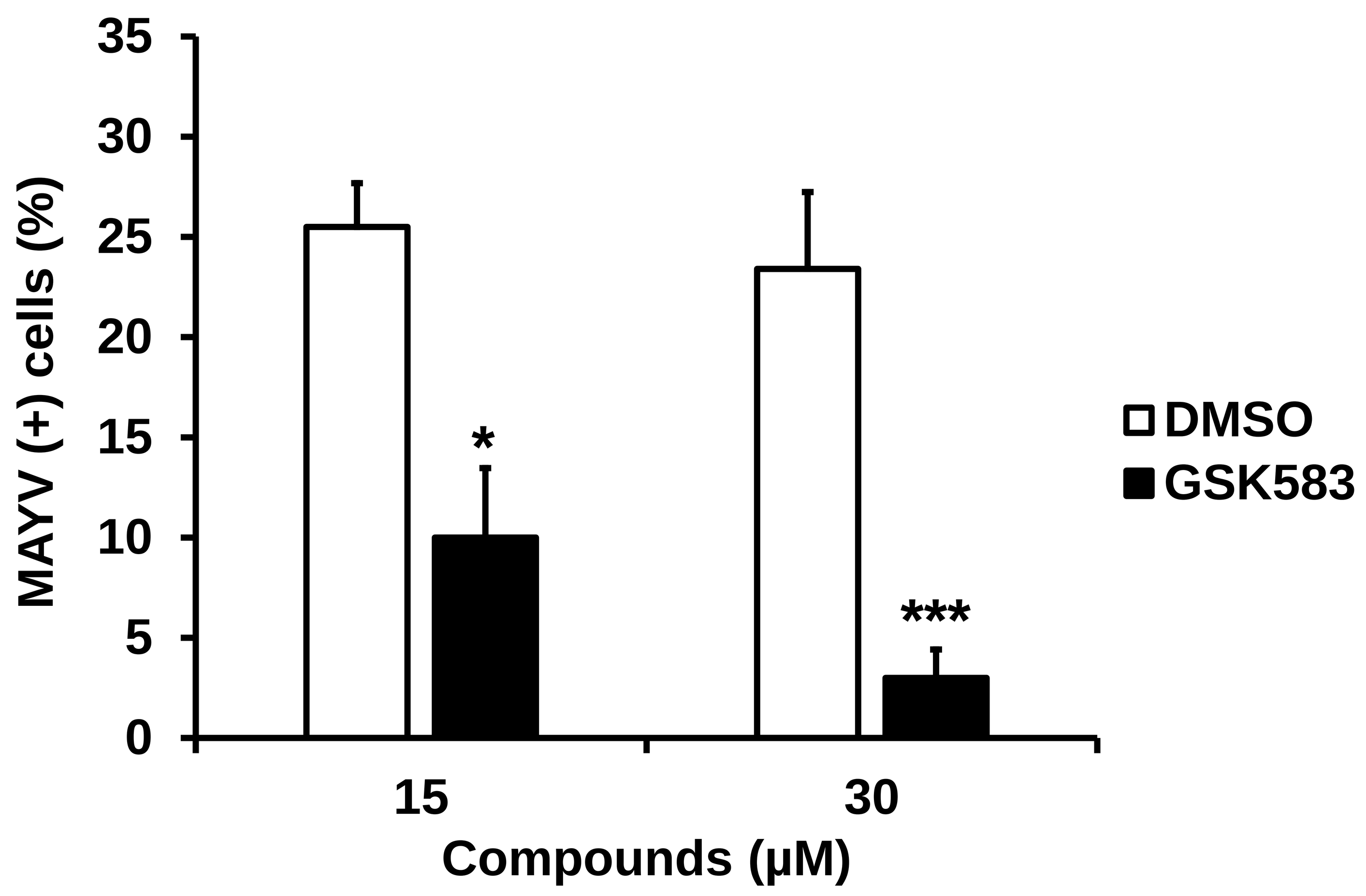
MAYV (+) cells (%)
35
30
25
20
15
10
5
0
*
***
15
30
Compounds (µM)
DMSO
GSK583

C

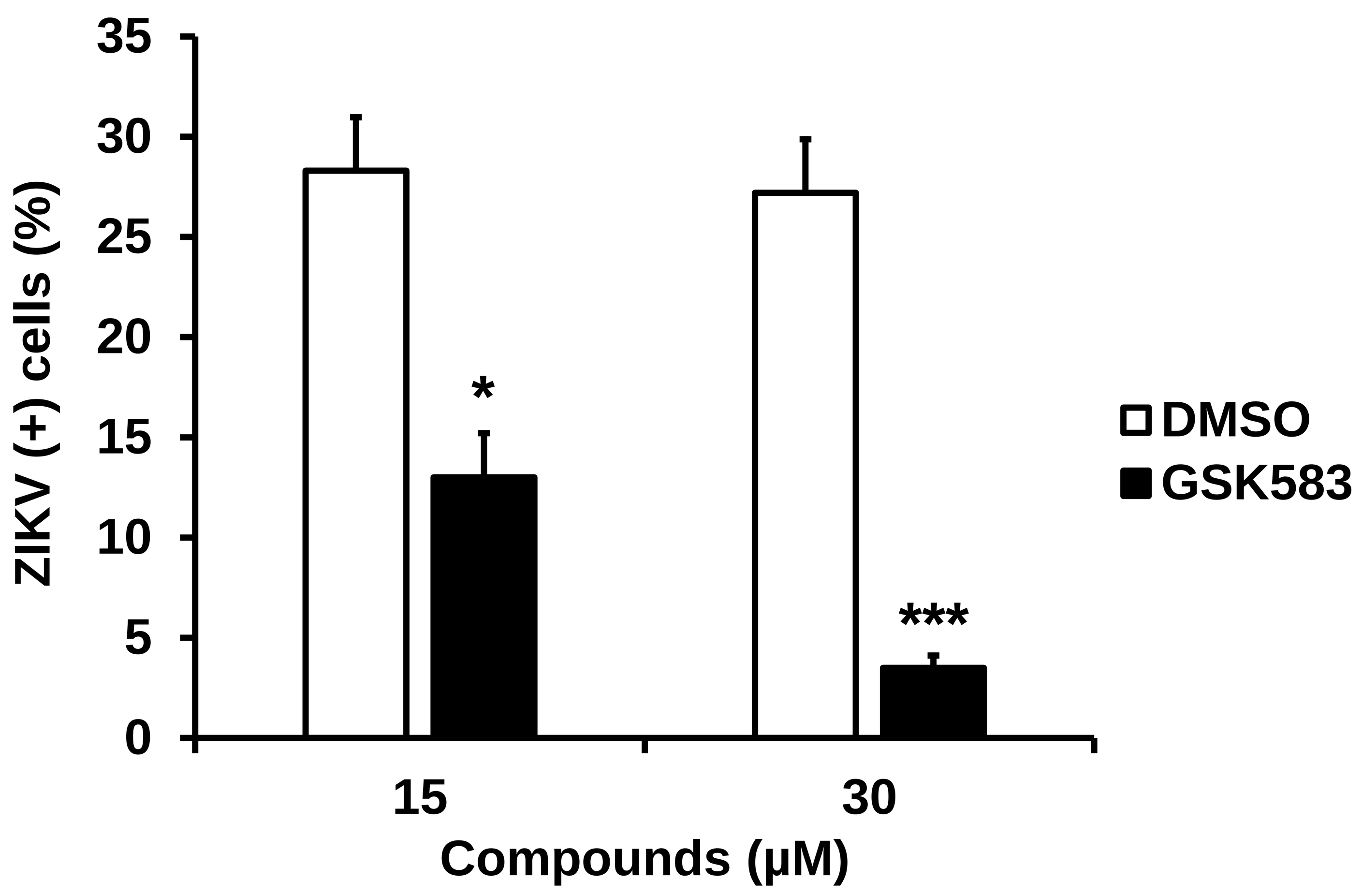
ZIKV (+) cells (%)
35
30
25
20
15
10
5
0
*
***
15
30
Compounds (µM)
DMSO
GSK583

D

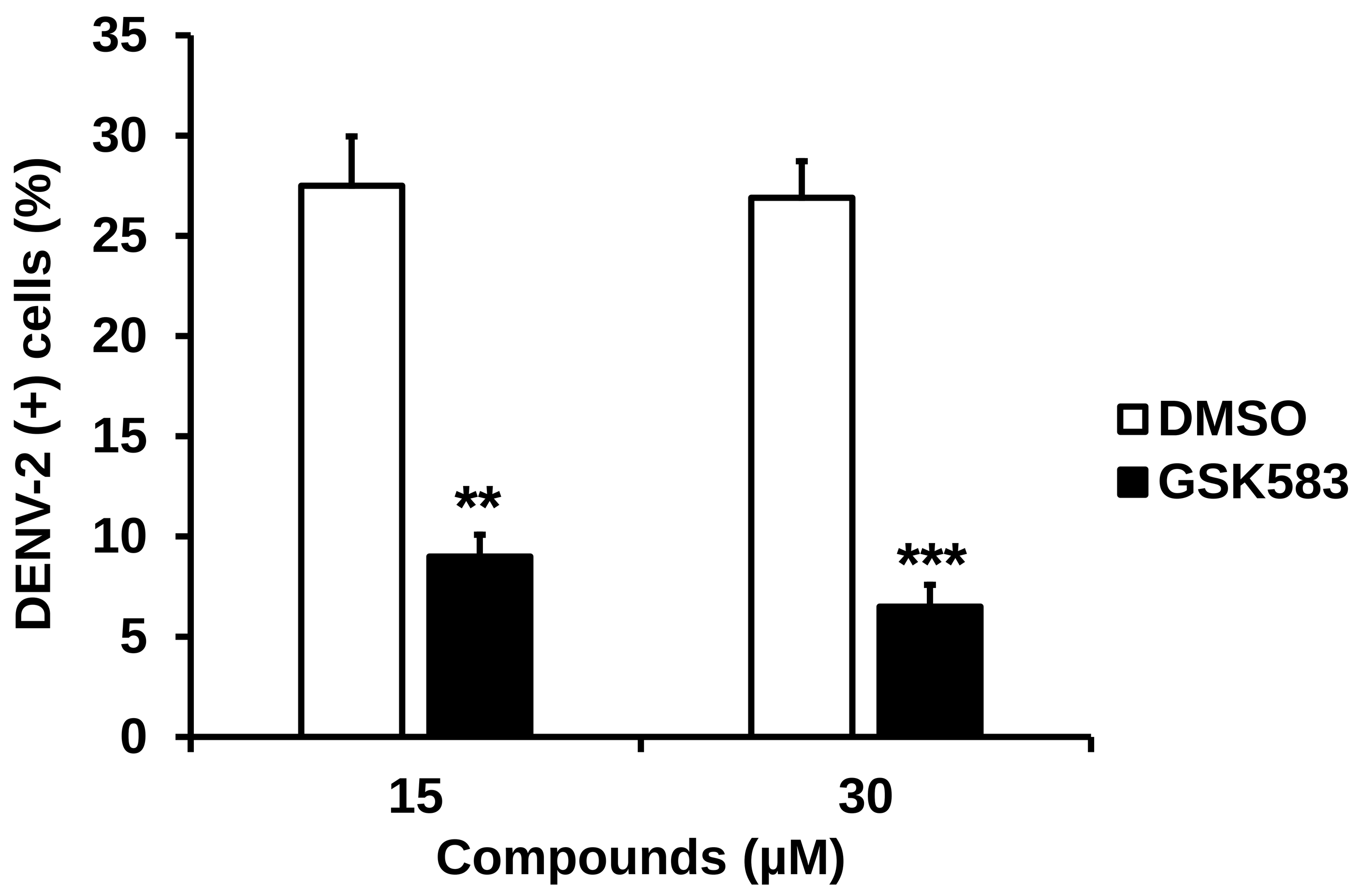
DENV-2 (+) cells (%)
35
30
25
20
15
10
5
0
**
***
15
30
Compounds (µM)
DMSO
GSK583

FIG S7

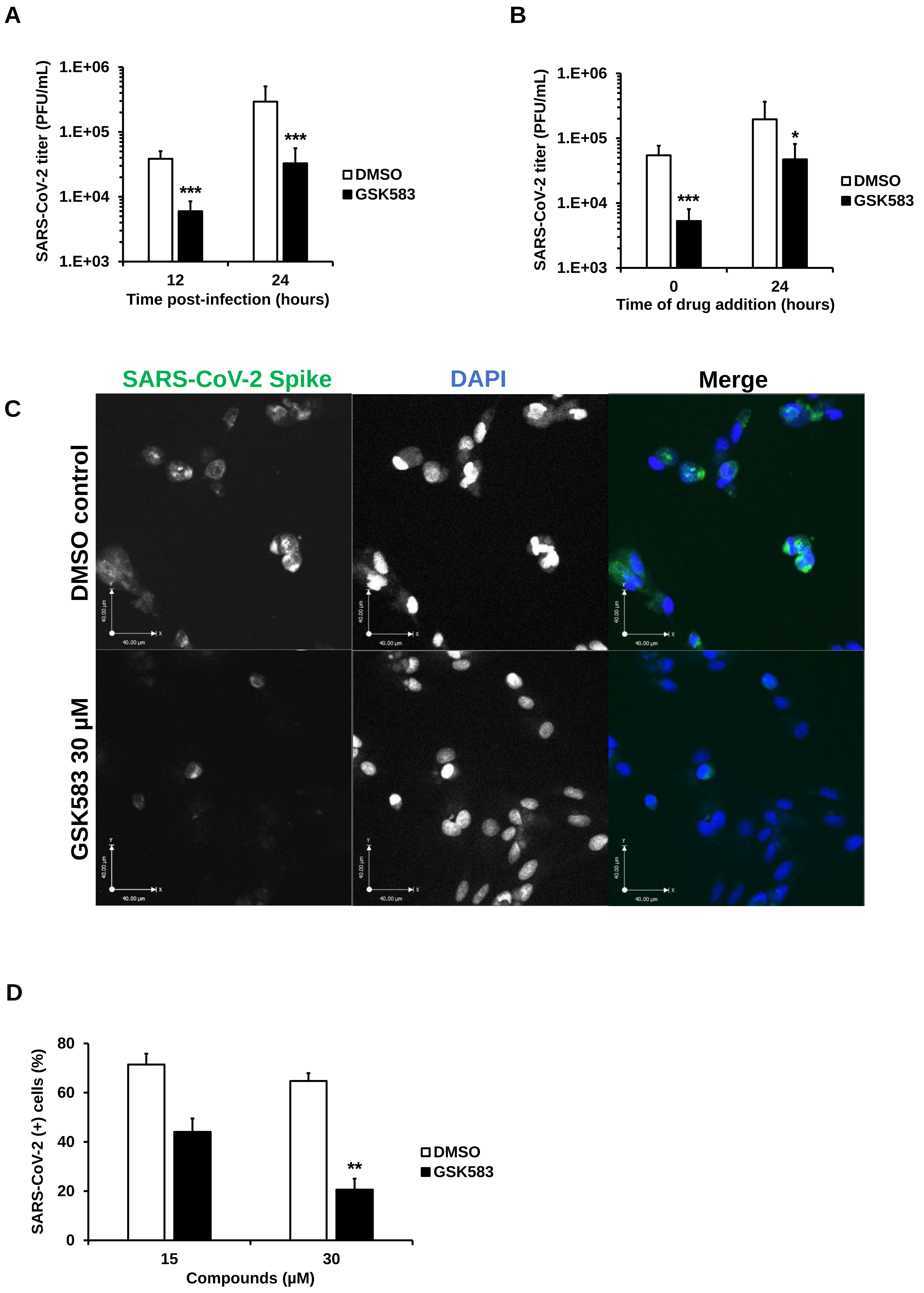
A
SARS-CoV-2 titer (PFU/mL)
1.E+06
1.E+05
1.E+04
1.E+03
***
***
12
24
Time post-infection (hours)
DMSO
GSK583
B
SARS-CoV-2 titer (PFU/mL)
1.E+06
1.E+05
1.E+04
1.E+03
***
*
0
24
Time of drug addition (hours)
DMSO
GSK583
C
SARS-CoV-2 Spike
DAPI
Merge
DMSO control
GSK583 30 μM
D
SARS-CoV-2 (+) cells (%)
80
60
40
20
0
**
15
30
Compounds (μM)
DMSO
GSK583

FIG S8

A

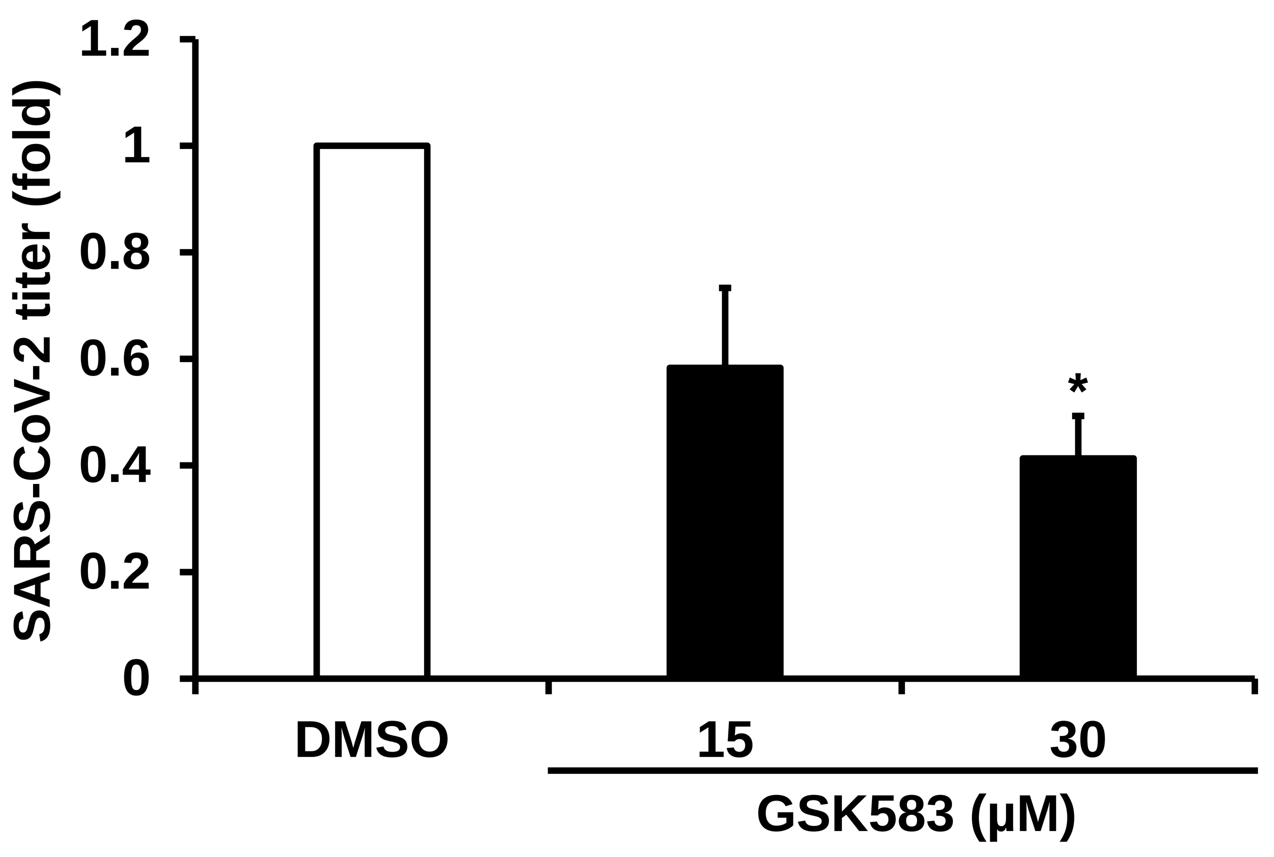

SARS-CoV-2 titer (fold)
1.2
1
0.8
0.6
0.4
0.2
0
*
DMSO
15
30
GSK583 (μM)


B

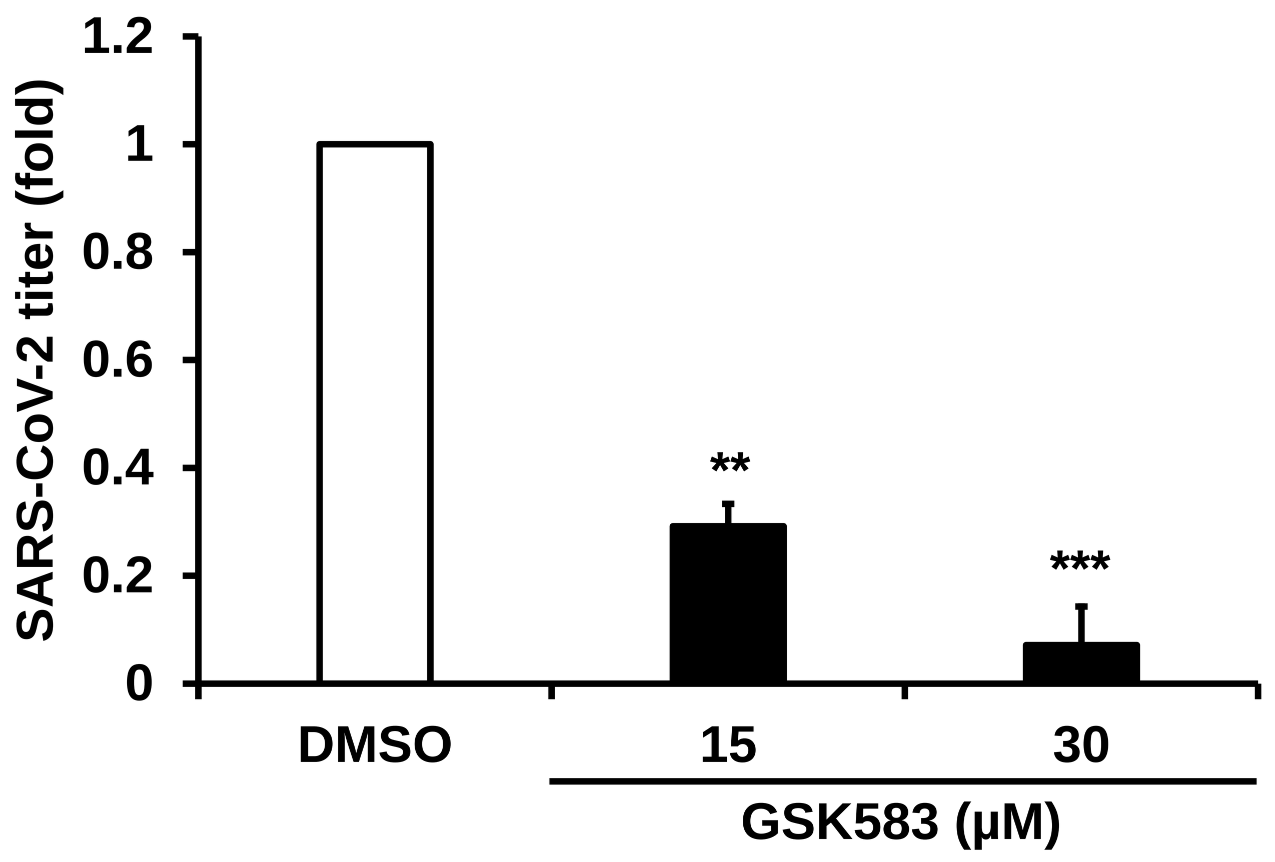

SARS-CoV-2 titer (fold)
1.2
1
0.8
0.6
0.4
0.2
0
**
***
DMSO
15
30
GSK583 (μM)


C

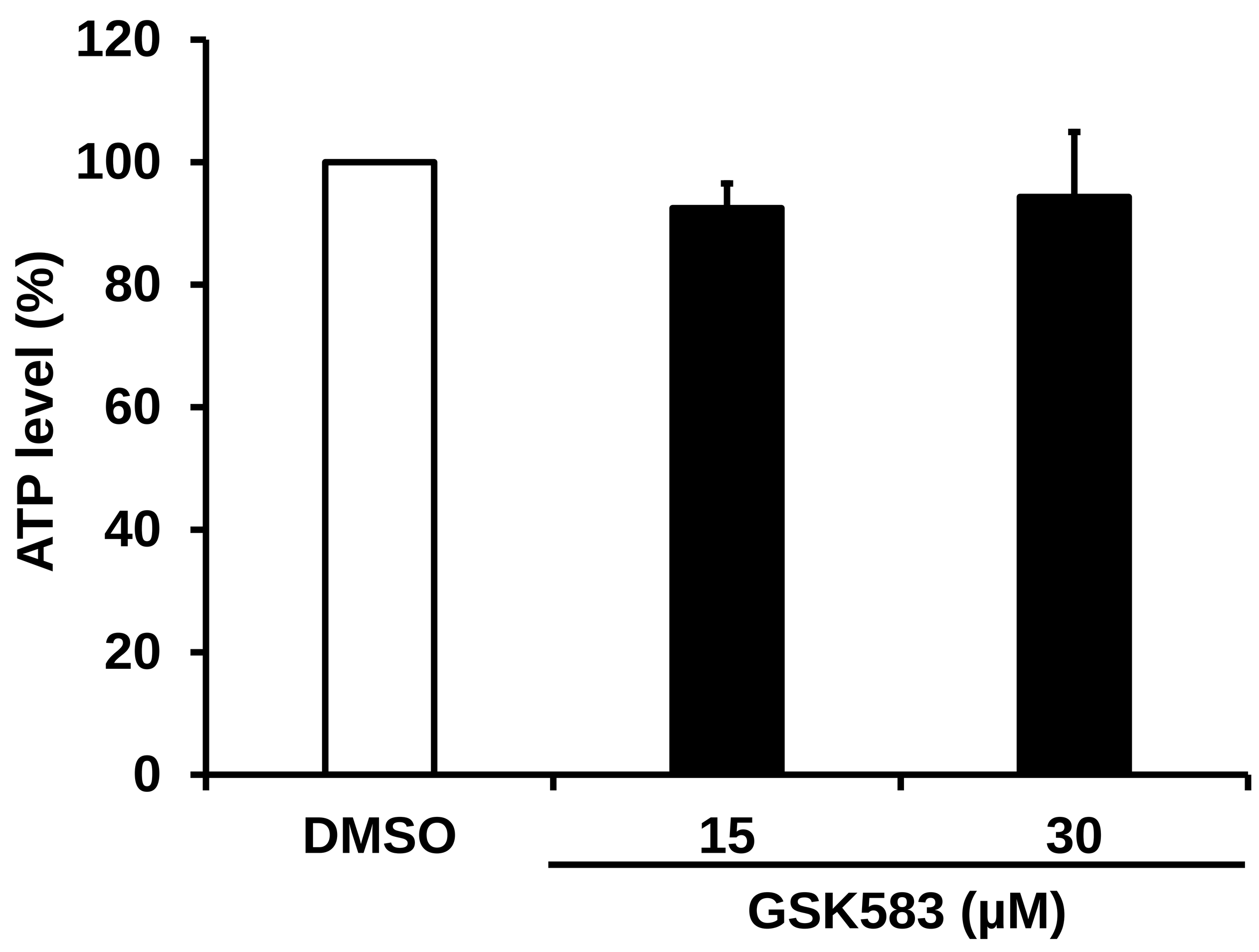

ATP level (%)
120
100
80
60
40
20
0
DMSO
15
30
GSK583 (μM)


D

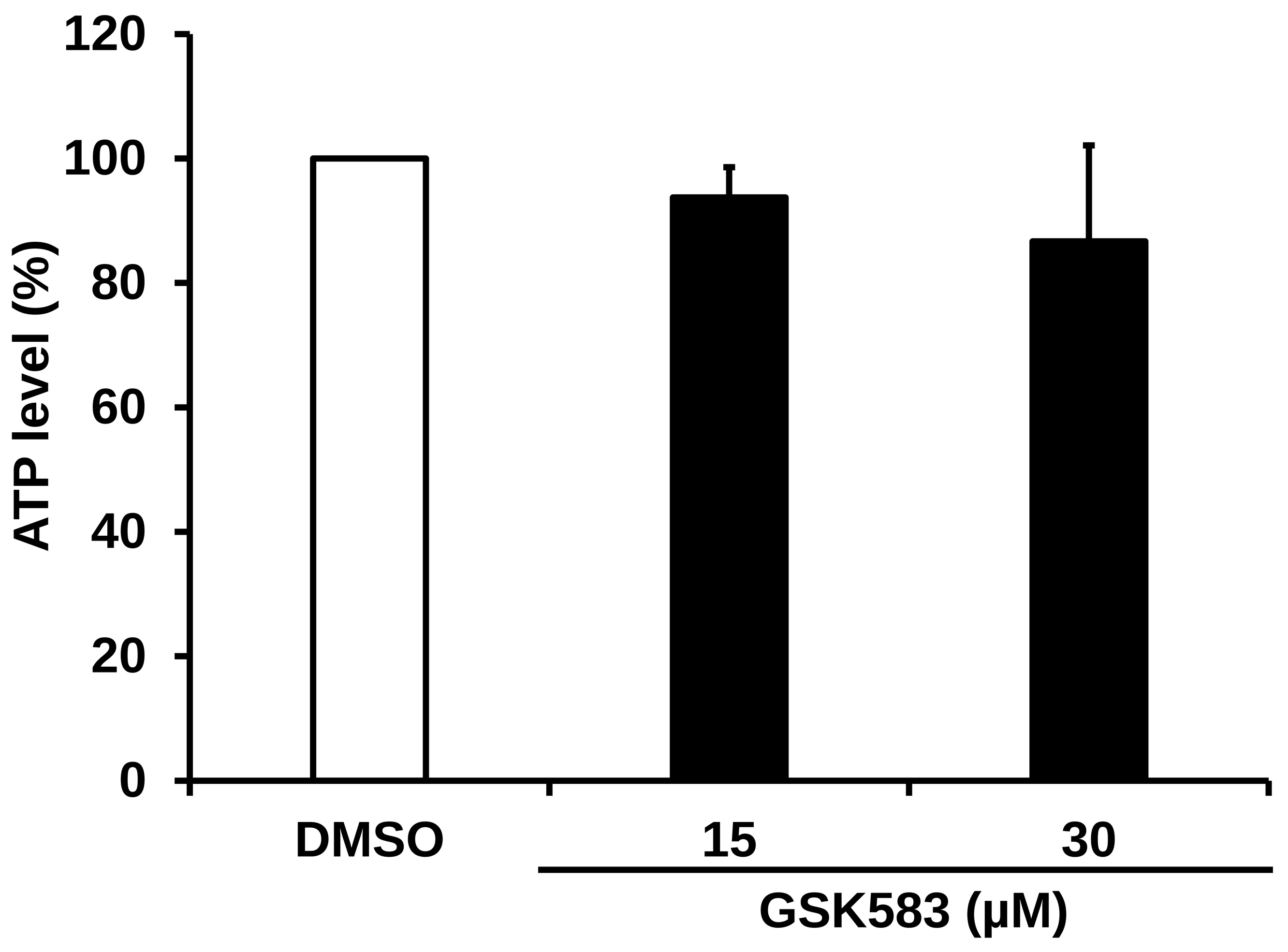

ATP level (%)
120
100
80
60
40
20
0
DMSO
15
30
GSK583 (μM)


E

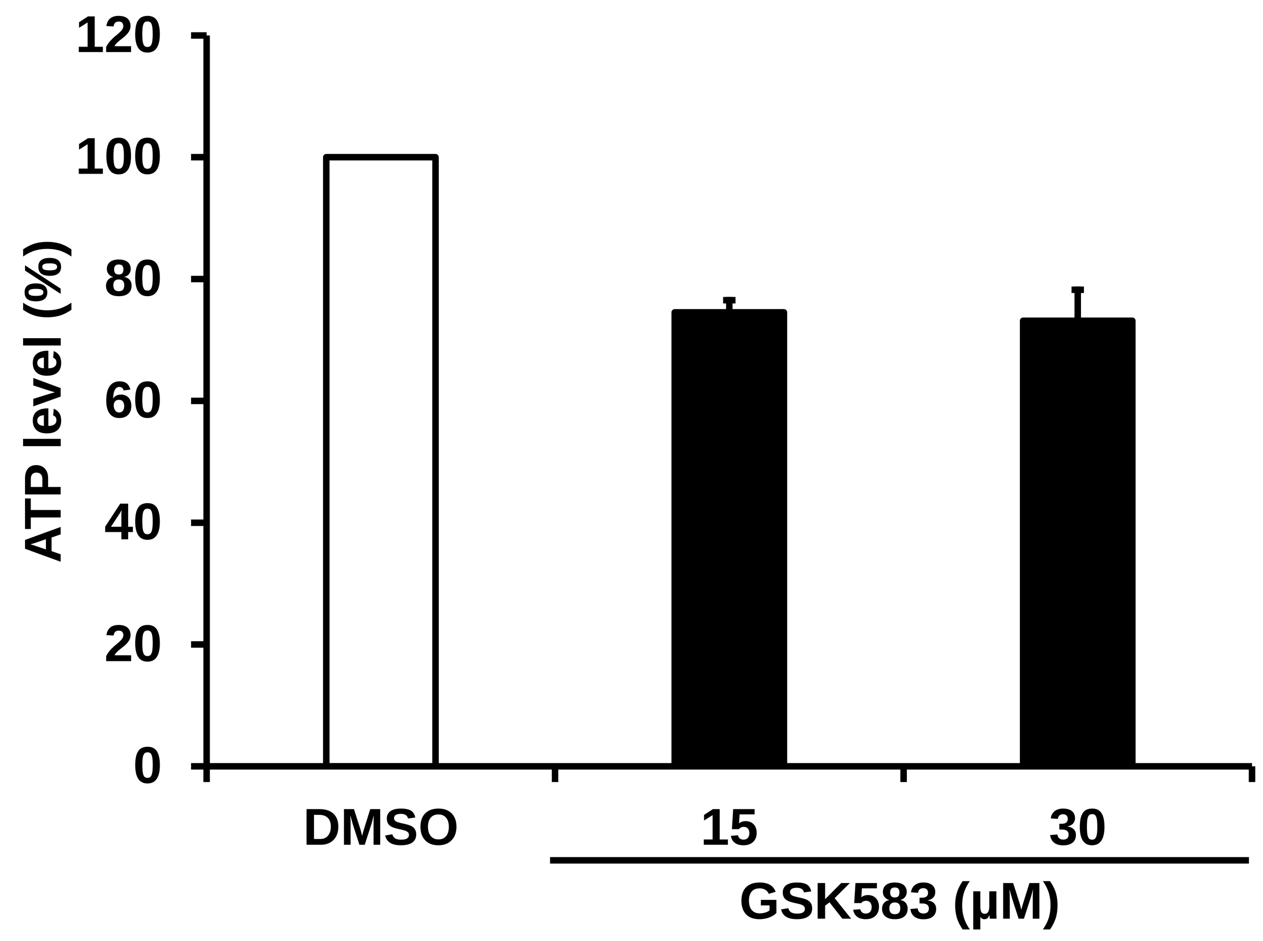

ATP level (%)
120
100
80
60
40
20
0
DMSO
15
30
GSK583 (μM)
